# Infection and chronic disease activate a brain-muscle signaling axis that regulates muscle performance

**DOI:** 10.1101/2020.12.20.423533

**Authors:** Shuo Yang, Meijie Tian, Yulong Dai, Shengyong Feng, Yunyun Wang, Deepak Chhangani, Tiffany Ou, Wenle Li, Ze Yang, Jennifer McAdow, Diego E. Rincon-Limas, Xin Yin, Wanbo Tai, Gong Cheng, Aaron Johnson

## Abstract

**Graphic abstract:** 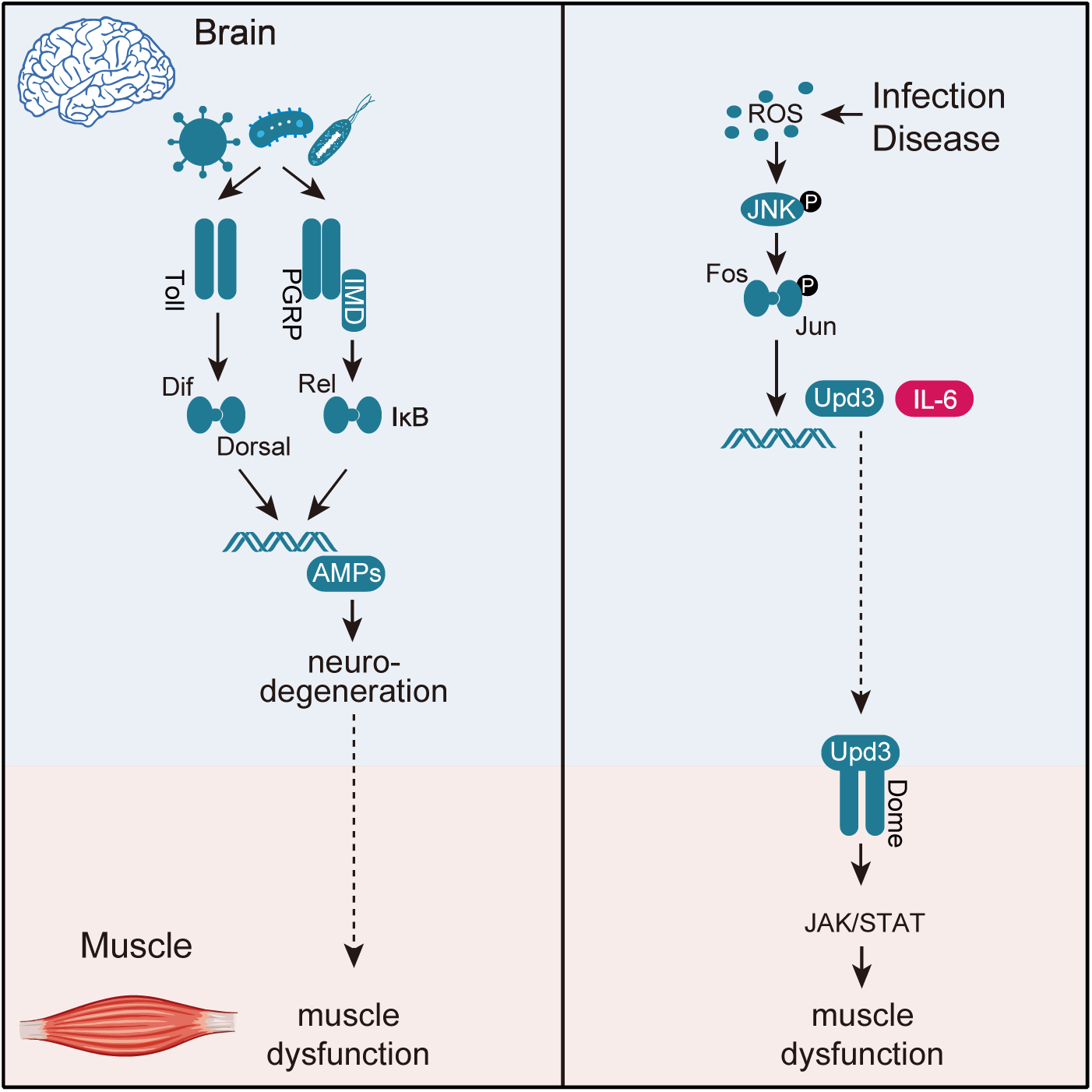

**Summary:** Infections and neurodegenerative diseases induce neuroinflammation, but affected individuals often show a number of non-neural symptoms including muscle pain and muscle fatigue. The molecular pathways by which neuroinflammation causes pathologies outside the central nervous system (CNS) are poorly understood, so we developed three models to investigate the impact of neuroinflammation on muscle performance. We found that bacterial infection, COVID-like viral infection, and expression of a neurotoxic protein associated with Alzheimer′ s disease promoted the accumulation of reactive oxygen species (ROS) in the brain. Excessive ROS induces the expression of the cytokine Unpaired 3 (Upd3) in insects, or its orthologue IL-6 in mammals, and CNS-derived Upd3/IL-6 activates the JAK/Stat pathway in skeletal muscle. In response to JAK/Stat signaling, mitochondrial function is impaired and muscle performance is reduced. Our work uncovers a brain-muscle signaling axis in which infections and chronic diseases induce cytokine-dependent changes in muscle performance, suggesting IL-6 could be a therapeutic target to treat muscle weakness caused by neuroinflammation.

## Introduction

Neuroinflammation refers to the activation of innate immune pathways in the central nervous system (CNS). Infectious diseases, including meningitis, Zika fever, and COVID-19, chronic conditions including Alzheimer′s disease and Parkinson’s Disease, and normal aging all induce neuroinflammation (Farmen et al., 2021; Frere et al., 2022; Hou et al., 2019; Leng and Edison, 2021; Lum et al., 2017). Although neuroinflammation can be activated by a number of factors, the inflammatory pathways appear to converge on a common disease mechanism that initiates neurodegeneration, defined as the disruption of neural function through changes in neuronal structure or survivability (Glass et al., 2010; Ransohoff, 2016). Symptoms associated with neurodegeneration are wide spread and include anxiety, insomnia, muscle weakness and even paralysis (Cao et al., 2013; Jayaraman et al., 2021; Leng and Edison, 2021). Neuroinflammation and neurodegeneration are thought to primarily target cells in the CNS, and symptoms in tissues outside of the CNS are due to changes in direct neural connectivity or function with a target tissue. However, neuroinflammation may also alter the secretome of the CNS, which could have profound effects on organ physiology outside of the CNS.

Inter-organ communication is emerging as a fundamental mechanism regulating whole-body physiology and homeostasis. Organs communicate using secreted molecules that enter circulation, translocate to target tissues, and then direct a variety of processes including immunity, behavior, neurogenesis, cardiovascular function, and cellular aging (Cai et al., 2021; Cao et al., 2022; Lee et al., 2014; Leiter et al., 2022; Robles-Murguia et al., 2020; Yang et al., 2019). Over 60 years ago it was proposed that skeletal muscle contractions liberate unknown factors that regulate metabolism (Goldstein, 1961). The discovery 40 years later that the cytokine IL-6 is released from skeletal muscle during exercise to regulate metabolism showed skeletal muscle is in fact an endocrine organ, which sparked great interest in understanding the role of muscle-derived signaling molecules, or myokines, on non-muscle physiology (Whitham and Febbraio, 2016). The CNS is a well-defined target of myokines, and over fifty muscle-derived proteins have been found to translocate from muscle to the brain (Droujinine et al., 2021). Myokines are now recognized as a heterogenous collection of proteins, that include conventional signaling molecules such as Bone Morphogenetic Proteins, as well as signaling accessory proteins such as transporters, that induce a variety of responses including the proliferation of neural precursors and the synthesis of neurotransmitters (Leiter et al., 2022; Robles-Murguia et al., 2020). Although the muscle-brain signaling axis has well defined roles in regulating neural activity, it remains unclear if a complementary brain-muscle signaling axis regulates muscle function.

IL-6 is a highly conserved extracellular ligand that activates the JAK/Stat pathway. JAK/Stat signaling fulfills a diverse array of developmental processes and homeostatic responses in insects and mammals, and the *Drosophila* genome encodes three IL-6 related ligands, Unpaired 1 (Upd1), Upd2, and Upd3 (Johnson et al., 2011; Moresi et al., 2019; Villarino et al., 2017). Upon binding to the receptor Domeless (Dome), Upd ligands activate the JAK/Stat pathway (Brown et al., 2001; Fisher et al., 2016). While JAK/Stat signaling is essential for development, regeneration, and immune responses, inappropriate JAK/Stat signaling activity can disrupt normal mitochondrial function in *Drosophila* skeletal muscle and in cultured mammalian muscle cells (Abid et al., 2020; Agaisse and Perrimon, 2004; Ding et al., 2021; Shen et al., 2022). In mice, IL-6 activated JAK/Stat signaling contributes to sepsis-induced muscle atrophy and weakness, and IL-6 serum levels in patients have been inversely correlated with functional muscle outcomes (Custodero et al., 2020; Grosicki et al., 2020). In addition, JAK/Stat inhibitors can improve muscle function in patients with Critical illness myopathy and other inflammation conditions (Addinsall et al., 2021; Chen et al., 2021b; Zanders et al., 2022). These data argue that tight regulation of JAK/Stat signaling is essential to maintain proper muscle function. Over the past several years the power of *Drosophila* to characterize inter-organ communication pathways (Rai et al., 2021; Robles-Murguia et al., 2020; Yang et al., 2019), and to identify pathogenic mechanisms driving infectious disease (Adamson et al., 2011; Chan et al., 2009; Hao et al., 2008; Harsh et al., 2020; Hughes et al., 2012; Liu et al., 2018; Yang et al., 2018) has come to light. We used *Drosophila* to investigate a putative brain-muscle signaling, and found neuroinflammation activates the brain-muscle axis and regulates muscle performance. By testing three models of neuroinflammation, that included *E. coli* infection in the brain, expression of a SARS-CoV-2 protein in the CNS, and expression of a neurotoxic amyloid-β protein associated with Alzheimer’s disease (AD) in the CNS, we uncovered a general mechanism in which Upd3 is expressed in the CNS in response to neuroinflammation, and subsequent JAK/Stat signaling in skeletal muscle disrupts mitochondrial function and reduces muscle performance. Incredibly, expressing SARS-CoV-2 proteins in the mouse brain also activated IL-6 expression, and reduced muscle performance. Long-COVID refers to a condition in which symptoms such as insomnia, muscle pain, and muscle fatigue persist after the SARS-CoV-2 virus is cleared from the respiratory tract (Subramanian et al., 2022; Xu et al., 2022). Bacterial infections in the CNS and AD are also associated with impaired muscle function (Beeri et al., 2021; Giannos et al., 2022; Martellosio et al., 2020). Our study argues neuroinflammation associated with bacterial infections, Long-COVID, and AD activates a conserved, IL-6 mediated brain-muscle signaling axis, suggesting IL-6 could be a therapeutic target for patients with infections and chronic diseases.

## Results

### Neural infection disrupts mitochondrial function in skeletal muscle

Neuroinflammation refers to the activation of innate immune pathways in the CNS, and bacterial infection in the *Drosophila* brain activates the innate immune response and impairs muscle function (Cao et al., 2013). To identify the pathways by which neuroinflammation affects muscle performance, we validated a neural infection model that uses direct infection of non-pathogenic *E. coli* to infect the fly brain and induce neuroinflammation. Infected animals had a significant reduction in climbing capacity compared to vehicle-injected controls at two- and six-days post-infection (Fig. 1A,B). The survival rate of infected flies was comparable to controls (Fig. S1A). Importantly, bacteria injected into the brain did not infect skeletal muscle (Fig. S1B), arguing the neural infection model induced neuroinflammation and impaired muscle performance without affecting survival or causing secondary infections in muscle. Muscle weakness could be caused by a number of factors, and we found flight muscle myofiber morphology was largely normal in infected animals but skeletal muscle mitochondrial activity was significantly reduced (Figs. 1C,D, S1C). Thus, infection-induced neuroinflammation disrupts mitochondrial function in skeletal muscle.

**Figure 1.**
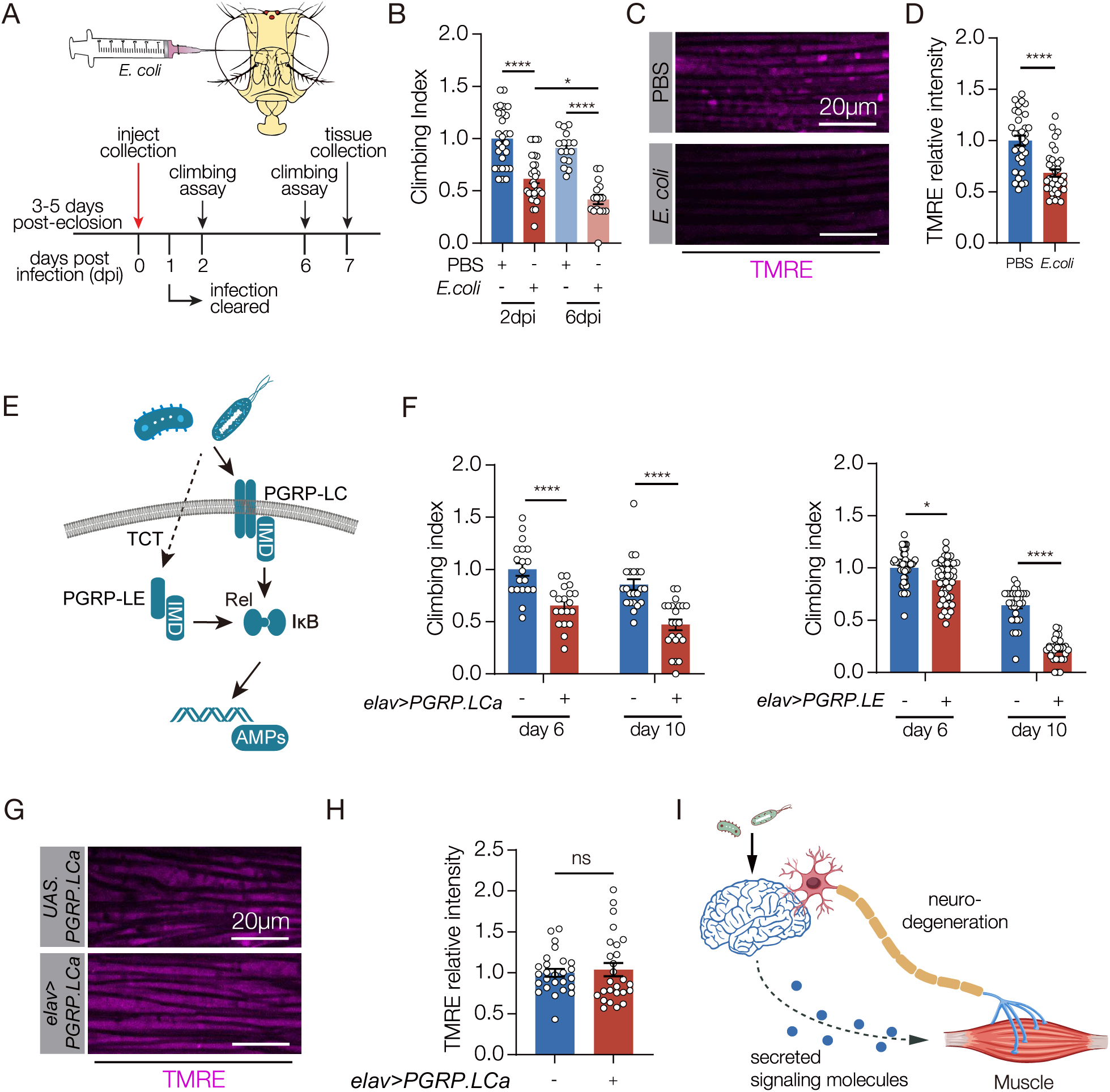
*E. coli* infection in the brain inhibits muscle performance. **A.** Experimental de-sign to study the skeletal muscle response to *E. coli* infection in the brain. **B.** Climbing index. The climbing capacity of sterile-injected (PBS, control) and *E. coli* injected flies was determined at 2- and 6-days post injury (dpi). Infected flies showed reduced climbing capacity. **C.** Confocal micrographs of indirect flight muscles stained with the potentiometric dye tetramethyl rhodamine ethyl ester (TMRE; violet) to assess mitochondrial membrane potential. Muscles from sterile injected flies showed more TMRE staining than *E. coli* injected flies. **D.** Quantification of the TMRE signal shown in C. **E.** Diagram of Drosophila Immune Deficiency (IMD) pathway. Infectious pathogens activate the Peptidoglycan receptor proteins (PGRPs), which activate Relish (Rel) and i-κB, that in turn initiate expression of antimicrobial peptides (AMPs) **F.** Climbing indexes. The climbing capacity of flies that expressed PGRPs in the CNS under the control of *elav.Gal4* (*elav>PGRPLCa, elav>PGRPLE*) was reduced at 6- and 10-days after eclosion compared to controls (*UAS.PGRPLCa, UAS.PGRPLE*). **G.** Confocal mi-crographs of indirect flight muscles stained with TMRE (violet). **H.** Quantification of the TMRE signal shown in G. TMRE levels were comparable between *elav>PGRPLCa* flies and controls (*UAS.PGRPLCa*). **I.** Model depicting two possible communication pathways between brain and muscle. Significance was determined by two-way ANOVA (B, F), and two-sided unpaired Student’s t-test (D, H). For climbing assays, data points represent the average of individual cohorts, with n≥15 flies per cohort. For TMRE, data points represent one of multiple fluorescent measurements per micrograph, with n≥5 flies per genotype. Error bars represent SEM. (*) p< 0.05, (**) p< 0.01, (***) p< 0.001, (****) p < 0.0001, (ns) non-significant.

To understand if neuroinflammation acts through the innate immune pathways to disrupt skeletal muscle function, we used our previously developed genetic model to activate the innate response in the CNS and assay muscle performance. In *Drosophila*, *E. coli* infection activates the Peptidoglycan receptor proteins (PGRPs) of the Immune Deficiency (IMD) pathway, which initiates the expression of antimicrobial peptides (AMPs) (Fig. 1E). AMPs in turn induce neurodegeneration, and inhibit climbing capacity (Cao et al., 2013). Transgenic expression of the PGRP-LC or the PGRP-LE receptor activates the IMD pathway in the absence of infection (Fig. S1D)(Yang et al., 2019), and we found flies that expressed PGRP-LC or PGRP-LE in the CNS showed significantly reduced climbing capacity compared to age matched controls (Fig. 1F). Similar to the bacterial infection model, myofiber morphology was unaffected in PGRP expressing flies, but surprisingly skeletal muscle mitochondrial activity was also unaffected (Figs. 1G,H, S1E). Bacterial infection therefore activates two pathways that impact muscle performance. An IMD-dependent pathway disrupts skeletal muscle performance but not mitochondrial activity, and an IMD-independent pathway regulates mitochondrial activity in skeletal muscle.

### Neural infection activates the brain-muscle signaling axis to regulate mitochondrial activity

Our studies of innate immune pathways showed inflammation-induced neurodegeneration does not regulate mitochondrial activity in skeletal muscle, raising the possibility that neuroinflammation activates a second mechanism that regulates muscle performance (Fig. 1I). Bacterial infection can alter the secretome of the infected tissue, and secreted proteins are often used for inter-organ communication (Cai et al., 2021). Infection outside the CNS induces *upd3* expression, and Upd3 is a secreted signaling ligand that activates the Janus kinase/signal transducer and activator of transcription (JAK/Stat) pathway (Sanchez Bosch et al., 2019). In addition, the JAK/Stat pathway regulates mitochondrial activity in *Drosophila* skeletal muscle and in cultured mammalian muscle cells (Abid et al., 2020; Ding et al., 2021). We hypothesized that bacteria-induced neuroinflammation releases Upd3 into circulation, which activates the JAK/Stat pathway in skeletal muscle and modulates mitochondrial function. Bacterial infection enriched *upd3* expression in the brain (Fig. 2A), and enhanced expression of the JAK/Stat target gene *socs36E* in skeletal muscle (Fig. 2A). However, *upd3* expression was unaffected in flies that expressed PGRP-LC or PGRP-LE in the CNS (Fig. S2A). The JAK/Stat pathway activates the transcription factor Stat92E, and *10XStat92E.GFP* is a validated reporter of Stat92E activity (Bach et al., 2007). *10XStat92E.GFP* expression in skeletal muscle was unaffected in flies that expressed PGRP-LC (Fig. S2B). These results argue the JAK/Stat pathway and innate immune pathways are independently activated in response to bacterial infection.

**Figure 2.**
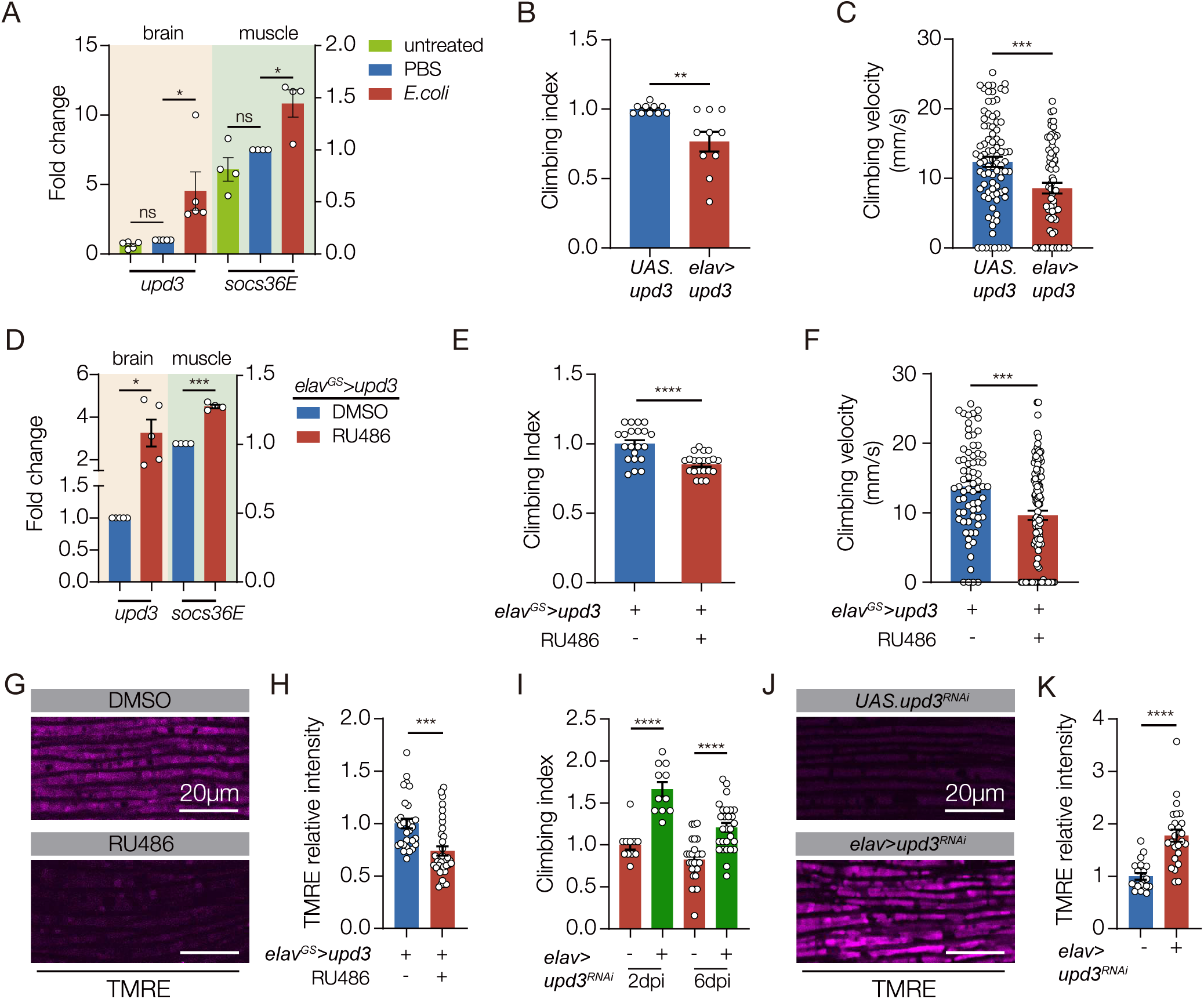
Neuroinflammation activates a Upd3 brain-muscle signaling axis. **A.** qRT-PCR. Untreated and sterile-injected control flies expressed less *upd3* in the brain and less *socs36e* in indirect flight muscles than *E. coli* injected flies at 7 days post injection (dpi). **B.** Climbing index. *elav>upd3* flies showed reduced climbing capacity compared to controls (*UAS.upd3*). **C.** Climbing velocity. *elav>upd3* flies climbed slower than controls. **D-G.** *elav^GS^.Gal4* was activated in adult flies with RU486 to induce *upd3* expression in the CNS. **D.** qRT-PCR. *elav^GS^>upd3* flies treated with RU486 showed more *upd3* in the brain and more *socs36e* in indirect flight muscles than flies treated with DMSO. **E.** Climbing index. *elav^GS^>upd3* flies treated with RU486 showed reduced climbing capacity compared to DMSO treated controls. **F.** Climbing velocity. *elav^GS^>upd3* flies treated with RU486 climbed slower than controls. **G.** Confocal micrographs of indirect flight muscles stained with TMRE (violet) to assess mitochondrial membrane potential. Muscles from *elav^GS^>upd3* flies treated with RU486 showed less TMRE staining than DMSO treated controls. H Quantification of the TMRE signal shown in G. **I.** Climbing index. *upd3^RNAi^* was used to knock down Upd3 expression in the CNS of *E. coli* infected flies. Infected flies with reduced Upd3 expression (*elav>upd3 ^RNAi^*) showed improved climbing capacity compared to controls at 2- and 6-dpi. **J.** Confocal micrographs of indirect flight muscles stained with TMRE (violet). Infected flies with reduced Upd3 expression (*elav>upd3 ^RNAi^*) showed improved mitochondrial membrane potential compared to controls at 6dpi. K. Quantification of the TMRE signal shown in J. Significance was determined by one-way ANOVA (A), two-sided unpaired student’s t-test (B-H), and two-way ANOVA (I). For qRT-PCR, data points represent biological replicates, with n≥5 flies per cohort. See Fig. 1 legend for Climbing Index and TMRE data points. Error bars represent SEM. (*) p< 0.05, (**) p< 0.01, (***) p< 0.001, (****) p < 0.0001, (ns) non-significant.

To understand if JAK/Stat signaling can modulate muscle performance, we overexpressed Upd3 in the CNS with *elav.Gal4*, and found *elav>upd3* flies had significantly reduced climbing ability (Fig. 2B,C). Since *elav.Gal4* is active in the embryo and can impede growth (Fig. S2C,D), we used the inducible gene-switch system (*elav^GS^.Gal4*) to activate *upd3* expression in the CNS of adult flies (hereafter, Upd3 gene switch). Upd3 gene switch flies showed enhanced *socs36E* expression in skeletal muscle, and reduced climbing capacity (Fig. 2D-F). Strikingly, mitochondrial activity in skeletal muscle was significantly reduced in Upd3 gene switch flies even though myofiber morphology was normal (Figs. 2G,H S2E).

These studies support the model that bacteria-induced neuroinflammation releases Upd3 into circulation, which in turn activates the JAK/Stat pathway in skeletal muscle to modulate mitochondrial function. To functionally test our model, we knocked down *upd3* in the CNS and found *upd3* knock down mitigated climbing defects associated with bacterial infection. Similarly, knocking down the Upd3 receptor *domeless* in skeletal muscle improved muscle performance in infected flies (Figs. 2I, S2F). *upd3* knock down in the CNS also improved muscle mitochondrial function in infected flies (Fig. 2J,K). Our data argue that bacterial infection activates a brain-muscle signaling axis in which CNS-derived Upd3 modulates mitochondrial activity in skeletal muscle.

### A viral model of neuroinflammation

Activating the brain-muscle axis could be a specific response that is limited to bacterial infection, or the brain-muscle axis could be activated in response to other pathogens that infect the CNS. To distinguish between these possibilities, we aimed to develop a clinically relevant model of viral infection in the CNS. A common symptom of COVID-19 is neuroinflammation, and the SARS-CoV-2 protein ORF3a activates the innate immune response (Crunfli et al., 2022; Frere et al., 2022; Song et al., 2021; Yang et al., 2021; Zhang et al., 2022). We asked if ORF3a protein is present in the brain of SARS-CoV-2 patients on autopsy, and, in the cerebellum, we detected ORF3a in the plasmalemma and cytolymph of Purkinje cells and in Granular cells of (Fig. 3A-C). ORF3a was also present in pyramidal cells of the hippocampus (Fig. 3C). ORF3a was not identified in samples from uninfected patients (Fig. 3C). Our studies argue ORF3a is neuroinvasive and could directly induce neuroinflammation, and align with recent observations showing SARS-CoV-2 can infect astrocytes in the brain and individual SARS-CoV-2 proteins, such as Spike protein, have been detected after the viral infection is cleared (Crunfli et al., 2022; Swank et al., 2022). ORF3a expression can therefore be used as a clinically relevant model of viral infection in the CNS.

**Figure 3.**
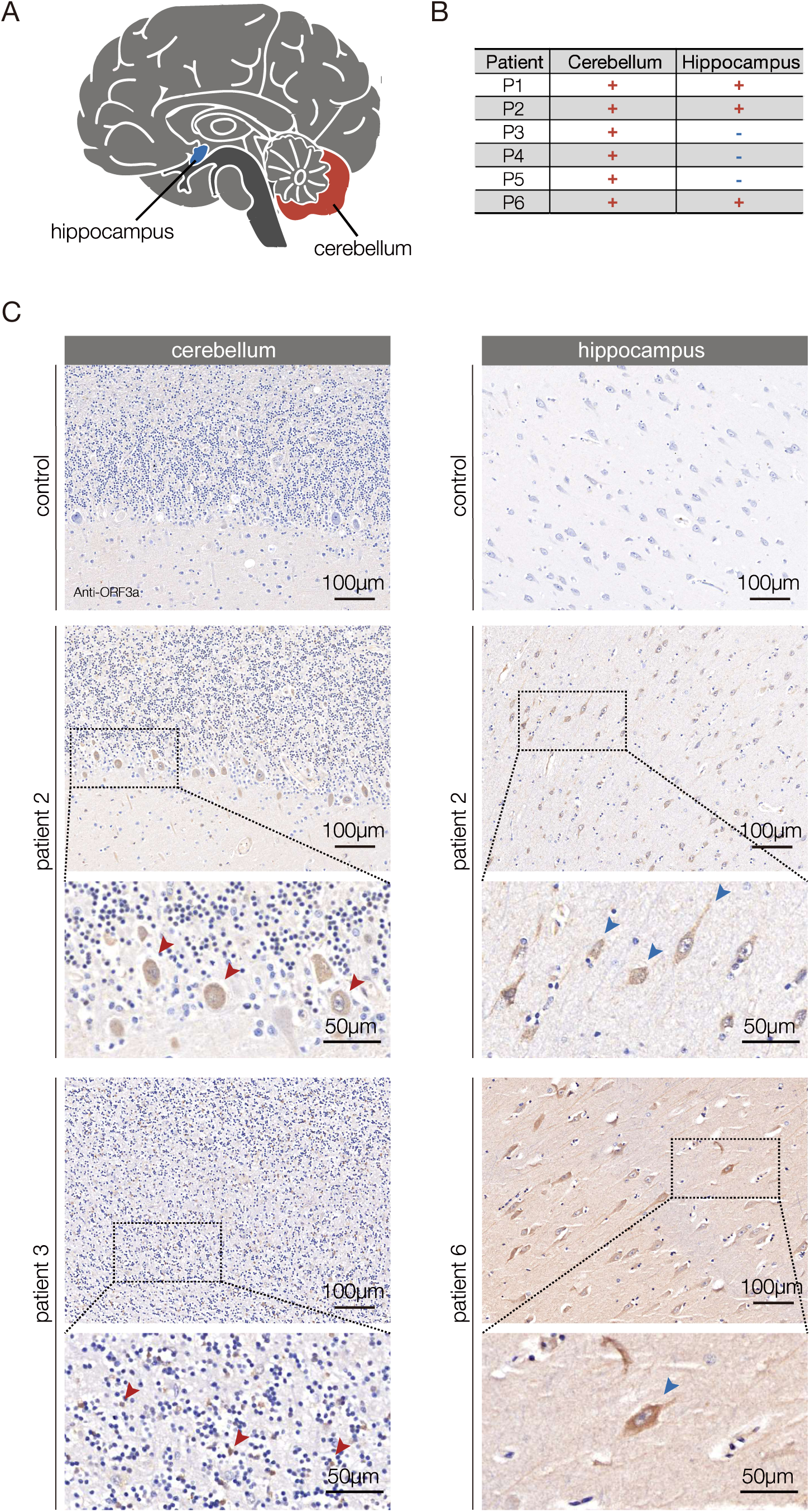
ORF3a is expressed in the brain of COVID-19 patients. **A.** Diagram of the human brain. **B.** Summary of the histological findings shown in C. The cerebellum and hippocampus samples from COVID-19 patients (patients 2,3 and 6) showed distinct and specific SARS-CoV-2 ORF3a antibody staining. Positive staining was not observed in uninfected controls. **C.** Micrographs of FFPE sections from COVID-19 patients and uninfected patients stained with an ORF3a antibody. In the cerebellum, ORF3a protein (brown) was detected in the plasmalemma and cytolymph of Purkinje cells (patient 2, red arrowheads) and Granular cells (patient 3, red arrows). In the hippocampus ORF3a protein was detected in pyramidal cells in (patients 2 and 6, blue arrowheads).

### ORF3a reduces muscle performance

We used the UAS-Gal4 system to express ORF3a in the *Drosophila* CNS (*elav>ORF3a)*, and found ORF3a expression alone was sufficient to activate neuroinflammation and reduce muscle performance (Fig. 4A-C). The life span of *elav>ORF3a* flies was also reduced (Fig. S3A). Similar to our bacterial infection model, skeletal muscle morphology was unaffected in *elav>ORF3a* flies (Fig. S3B). Interestingly, muscle performance and longevity were not affected in flies that expressed ORF3a in muscle, suggesting ORF3a acts cell non-autonomously to regulate muscle physiology (Fig. S3C,D).

**Figure 4.**
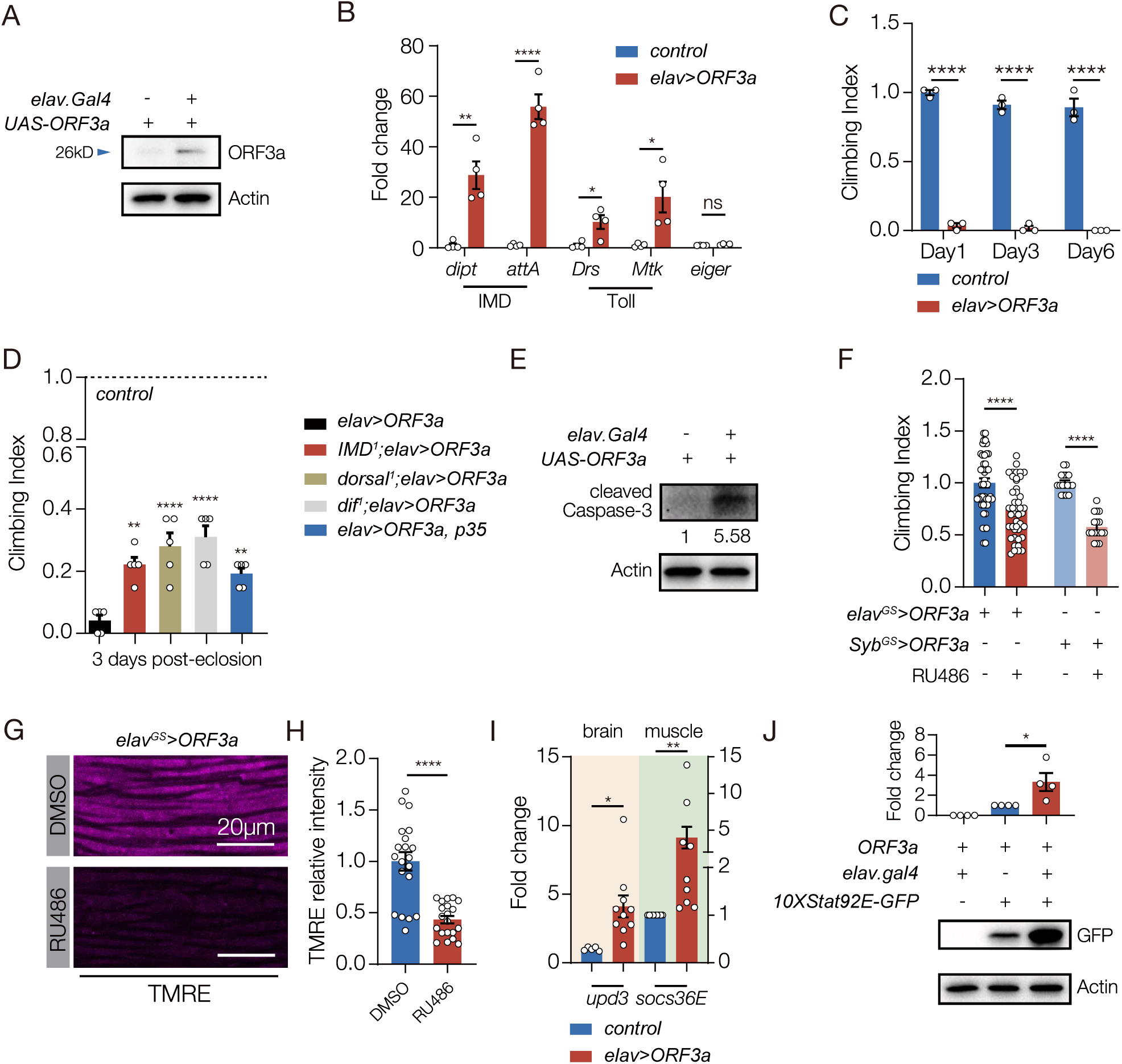
ORF3a induces neuroinflammation and reduces muscle performance in *Drosophila.* **A.** Western blot. ORF3a protein was expressed in the brain of *elav>ORF3a* flies but not in control flies (*UAS.ORF3a*). ORF3a was detected at ∼26kD. **B.** qRT-PCR. Transcripts encoding the antimicrobial peptides (AMPs) activated by the IMD pathway (*Dipt* and *attA*) and the Toll pathway (*Drs* and *Mtk*) were enriched in the brain of *elav>ORF3a* flies compared to controls (*UAS.ORF3a*). The cytokine *eiger* is a TNF orthologue that can induce apoptosis in response to infection (Igaki and Miura, 2014), but *eiger* expression was unchanged in *elav>ORF3a* flies. **C.** Climbing index. *elav>ORF3a* flies showed reduced climbing capacity compared to controls (*UAS.ORF3a*) at 1-, 3-, and 6-days after eclosion. **D.** Climbing index. *elav>ORF3a* flies with homozygous mutations affecting the IMD pathway (*IMD^1^*) or the Toll pathway (*dorsal^1^*, *dif^1^*) showed improved climbing capacity compared to *elav>ORF3a* flies. *elav>ORF3a* flies that expressed the inhibitor of apoptosis p35 in the CNS also showed im-proved climbing capacity compared to *elav>ORF3a* flies. Control flies (*UAS.ORF3a*) were used to establish the base of the index. **E.** Western blot. Cleaved Caspase-3 was enriched in the brain of *elav>ORF3a* flies compared to controls (*UAS.ORF3a*). **F,G.** *elav^GS^.Gal4* and *Syb^GS^.Gal4* were activated in adult flies with RU486 to induce *ORF3a* expression in the CNS. **F.** Climbing index. *elav^GS^>ORF3a* and *Syb^GS^.ORF3a* flies treated with RU486 showed re-duced climbing capacity compared to DMSO treated controls. **G.** Confocal micrographs of indirect flight muscles stained with TMRE (violet) to assess mitochondrial membrane poten-tial. Muscles from *elav^GS^>ORF3a* flies treated with RU486 showed less TMRE staining than DMSO treated controls. H. Quantification of the TMRE signal shown in G. **I.** qRT-PCR. *elav>ORF3a* flies expressed more *upd3* in the brain and more *socs36e* in indirect flight mus-cles than control flies (*UAS.ORF3a*) at 7 days after eclosion. **J.** Western blot. GFP expression from the JAK/Stat activity reporter *10XStat92E.GFP* in muscle was enriched in *elav>ORF3a* flies compared to controls (*UAS.ORF3a*). Significance was determined by two-sided unpaired Student’s t-test (F, H, I, J), two-way ANOVA (B, C), and one-way ANOVA (D). For Western blots and qRT-PCR, relative expression was determined for three biological replicates. See Fig. 1 legend for Climbing Index data points. Error bars represent SEM. (*) p< 0.05, (**) p< 0.01, (***) p< 0.001, (****) p < 0.0001, (ns) non-significant.

### ORF3a causes Long-COVID like symptoms

Long-COVID refers to ongoing clinical symptoms, including memory loss, anxiety, insomnia, muscle weakness, and fatigue, after SARS-CoV-2 infection is no longer detectable in the respiratory tract (Huang et al., 2021; Taquet et al., 2021). Persistent neuroinflammation is observed in post-infectious patients and might be a major cause of Long-COVID. ORF3a expressing flies showed neuroinflammation, muscle weakness, and fatigue and we asked if ORF3a expression caused any additional Long-COVID like symptoms. Insomnia is a symptom associated with Long-COVID (Thompson et al., 2022), and locomotor activity can be used to study circadian rhythms and sleep parameters in *Drosophila* (Chiu et al., 2010; Seugnet et al., 2009). Flies that expressed ORF3a in the CNS showed insomnia-like phenotype at night, but animals that expressed ORF3a in glia showed normal circadian behavior (Fig. S3E-H). These data highlight the possibility that Long-COVID like symptoms, including neuroinflammation, insomnia, and muscle weakness, could be caused by the persisting presence of ORF3a, and other SARS-CoV-2 proteins, after the SARS-CoV-2 virus has been cleared. In addition, our studies argue ORF3a expression can used to model Long-COVID like symptoms.

### ORF3a activates innate immune pathways and the brain-muscle signaling axis

The mechanism by which ORF3a affects skeletal muscle performance could mimic that of *E. coli* infection in which neuroinflammation initiates AMP-mediated neurodegeneration via the innate immune pathways, and concurrently activates the Upd3-mediated brain-muscle signaling axis. We used enhancer-suppressor studies to test the possibility that ORF3a-mediated activation of the innate immune pathways regulates muscle performance and longevity, and found mutations in components of the IMD and the Toll pathways suppressed the *elav>ORF3a* phenotypes (Fig. 4D). The Toll and IMD pathways can also initiate AMP-mediated cell death (Cao et al., 2013), and flies that expressed ORF3a in the CNS showed elevated apoptosis in the brain (Fig. 4E). We inhibited apoptosis in *elav>ORF3a* flies by co-expressing p35, and found muscle performance was improved and life span was extended (Figs. 4D, Fig. S3I). These data suggest ORF3a activates an innate immune response that promotes neurodegeneration and reduces muscle performance.

To understand if ORF3a also activates the Upd3-mediated brain-muscle signaling axis, we used the inducible gene-switch system to express ORF3a in the adult CNS, and found ORF3a gene switch flies had reduced climbing capacity (Fig. 4F). Importantly, ORF3a gene switch flies also showed reduced mitochondrial activity in skeletal muscle, which is a phenotype specific to Upd3 overexpression (Fig. 4G,H). Flies that expressed ORF3a in the CNS showed enriched *upd3* expression in the brain, and enriched *socs36E* expression in skeletal muscle (Fig. 4I). Furthermore, *elav>ORF3a* flies showed a dramatic induction of *10XStat92E.GFP* expression in skeletal muscle, arguing ORF3a expression in the CNS directs a Stat92E transcriptional response in skeletal muscle (Fig. 4J). ORF3a therefore regulates skeletal muscle performance by activating Upd3-mediated brain-muscle signaling axis, and by initiating an innate immune response that promotes neurodegeneration.

### The brain-muscle axis is activated by ROS

How then does persistent ORF3a expression activate the brain-muscle axis and induce Long-COVID like symptoms? ROS production can induce Upd3/IL-6 expression across species (Gera et al., 2022; Henriquez-Olguin et al., 2015; Santabarbara-Ruiz et al., 2015), so we asked if ORF3a could also activate ROS production. HEK293 and HeLa cells transfected with ORF3a showed significantly higher ROS levels than control transfected cells (Figs. S4A, B), and *elav>ORF3a* flies had elevated ROS levels in the brain (Fig.5A). In addition, *elav>ORF3a* flies treated with a ROS scavenger that reduces ROS levels showed improved muscle performance compared to untreated controls (Fig.5B). ROS activates the JNK pathway to induce Upd3 expression (Gera et al., 2022), and the levels of phosphorylated JNK were elevated in *elav>ORF3a* flies (Fig.5C). Importantly, ROS production was unaffected in *elav>upd3* flies, arguing Upd3 acts downstream of an ORF3a-ROS-JNK signaling cascade (Figs.5D).

**Figure 5.**
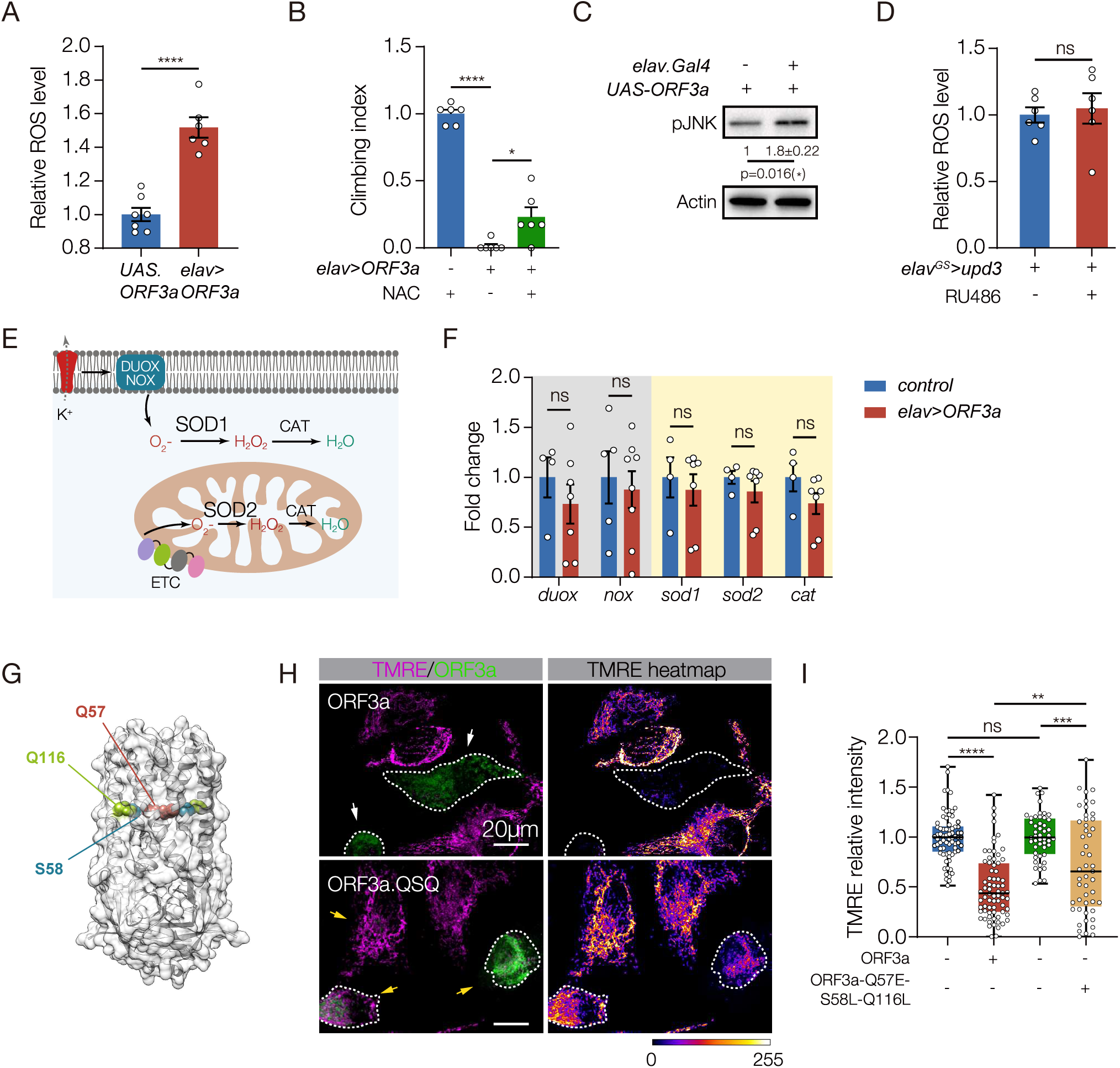
ORF3a enhances ROS in the brain and reduces mitochondrial function in skeletal muscle. **A.** ROS assay. H2DCFDA was used to measure ROS in the brain. *elav>ORF3a* flies produced more ROS than controls (*UAS.ORF3a*). **B.** Climbing index. *elav>ORF3a* flies treated with the ROS inhibitor N-acetyl-L-cysteine (NAC) showed improved climbing capacity compared to vehicle-treated controls. Wild-type flies treated with NAC were used to establish the base of the index. **C.** Western blot. pJNK levels in the brain were enriched in *elav>ORF3a* flies compared to controls (*UAS.ORF3a*). **D.** ROS assay. *elav^GS^>upd3* flies treated with RU486 accumulated more ROS in the brain than DMSO treated controls. **E.** Diagram of canonical ROS pathways. **F.** qRT-PCR. The expression of ROS-related transcripts in the brain was comparable between *elav>ORF3a* and control (*UAS.ORF3a*) flies. **G.** Cryo-EM structure of the ORF3a dimer, modified from (Kern et al., 2021). Residues positioned at the top of the polar cavity (Q57) or at the base of the hydrophilic grooves (S58, Q116) are shown. **H.** Micrographs of HeLA cells transfected with eGFP.ORF3a or triple mutant eGFP.ORF3a.QSQ (Q57E, S58L, Q116L), labeled for GFP (green) and TMRE (violet) to measure mitochondrial membrane potential. Cells transfected with wild-type ORF3a showed less TMRE fluorescence than cells transfected with ORF3a.QSQ. Dotted lines outline transfected cells. **I.** Quantification of TMRE fluorescence shown in H. Significance was determined by two-sided unpaired student’s t-test (A, C, D, F), and one-way ANOVA (B, I). For Western blots, qRT-PCR, and ROS assays relative expression was determined for a minimum of three biological replicates. See Fig. 1 legend for Climbing Index data points. Data points for TMRE expression represent fluorescence in a single transfected cell normalized to untransfected cells within the same field. n=10 fields. Error bars represent SEM. (*) p< 0.05, (**) p< 0.01, (***) p< 0.001, (****) p < 0.0001, (ns) non-significant.

Members of the nicotinamide adenine dinucleotide phosphate (NADPH) oxidase (NOX) and dual oxidase (DUOX) family of proteins are the primary producers of cellular ROS and, in response to bacterial and viral infections, NOX and DUOX derived ROS contribute to inflammation and tissue damage (Khomich et al., 2018; Lee et al., 2015). The mitochondrial electronic transport chain (ETC) also generates ROS through a NOX and DUOX independent mechanism (Zhao et al., 2019). The super oxidase dismutase (SOD) and catalase (CAT) enzymes reduce ROS levels and detoxify the cellular environment in response to ROS production (Fig.5E). To understand how SARS-CoV-2 infections might alter ROS levels, we assayed *nox* and *duox* expression in the brain, but found *nox* and *duox* expression was unaffected in *elav>ORF3a* flies (Fig.5F). A second possibility is that ORF3a reduces SOD and CAT levels, which would increase ROS levels. However, the expression of *sod1, sod2,* and *cat* in the brain was also unaffected in *elav>ORF3a* flies (Fig.5F).

We next considered the possibility that ORF3a activates ROS production through an atypical mechanism. Viroporins are hydrophobic proteins encoded by some viruses that aggregate in host cell membranes (Nieva et al., 2012). The influenza virus M2 protein is a well-studied viroporin, and M2 functions as an ion channel that induces mitochondrial dysfunction and ROS production (Moriyama et al., 2019). ORF3a is also a viroporin, and functions as a nonselective, Ca^2+^ permeable ion channel (Kern et al., 2021). Missense mutations in residues positioned at the top of the polar cavity (Q57E) or at the of the base of the hydrophilic grooves (S58L, Q116L) partially reduced ORF3a ion channel activity (Fig.5G) (Kern et al., 2021). In addition, ORF3a proteins containing either the Q57E mutation or the S58L, Q116L mutations partially attenuated ORF3a-induced cellular phenotypes (Chen et al., 2021a). We hypothesized that ORF3a activates ROS production through its viroporin activity, and generated a triple mutant that disrupts the polar cavity and the hydrophilic grooves (Q57E/S58L/Q116L; hereafter ORF3a.QSQ). Cells transfected with wild-type ORF3a showed a dramatic reduction in mitochondrial activity, whereas mitochondrial activity in cells transfected with ORF3a.QSQ was largely normal (Fig.5H,I; Fig. S4C). These data argue ORF3a ion channel activity contributes to mitochondrial pathogenicity, and suggest ORF3a induces ROS production by disrupting ion homeostasis. ORF3a expression in the CNS therefore regulates muscle performance by activating a ROS-JNK-Upd3-JAK/Stat signaling pathway that impairs mitochondrial function in skeletal muscle.

### ORF3a affects muscle performance in mammals

We wanted to understand if ORF3a could also induce neuroinflammation and disrupt muscle performance in mammals. Although ORF3a induced IL-1β and IL-8 expression through an NF-κB-dependent mechanism in human cell lines (Gowda et al., 2021), the pathogenicity of ORF3a in mammals has not been assessed *in vivo*. Adeno-associated virus (AAV) strategies to deliver SARS-CoV-2 coding sequences successfully showed the N protein promotes inflammation and can induce lung injuries in mice (Pan et al., 2021). We used a similar strategy involving retro-orbital AAV injections to deliver ORF3a to the frontal cortex of adult mice (Figs. 6A, S5A-C). ORF3a activated the expression of multiple cytokines, including IL-1β, CXCL-15, and the mammalian orthologue of Upd3, IL-6 (Fig. 6B,C). ORF3a expressing mice also showed a transient reduction in bodyweight (Fig. 6D). Similar to our results in *Drosophila*, ORF3a induced apoptosis and enhanced ROS production in the mammalian CNS (Fig. 6E,F). Strikingly, ORF3a expressing mice showed significant fatigue during treadmill running from four to sixteen days after AAV injection (Fig. 6G). In addition, ROS levels were elevated in skeletal muscles of ORF3a-expressing mice, suggesting mitochondrial dysfunction impaired muscle function (Fig. 6H). These results argue ORF3a expression in mice disrupts muscle performance, and suggest neuroinflammation activates the brain-muscle axis in mammals.

**Figure 6.**
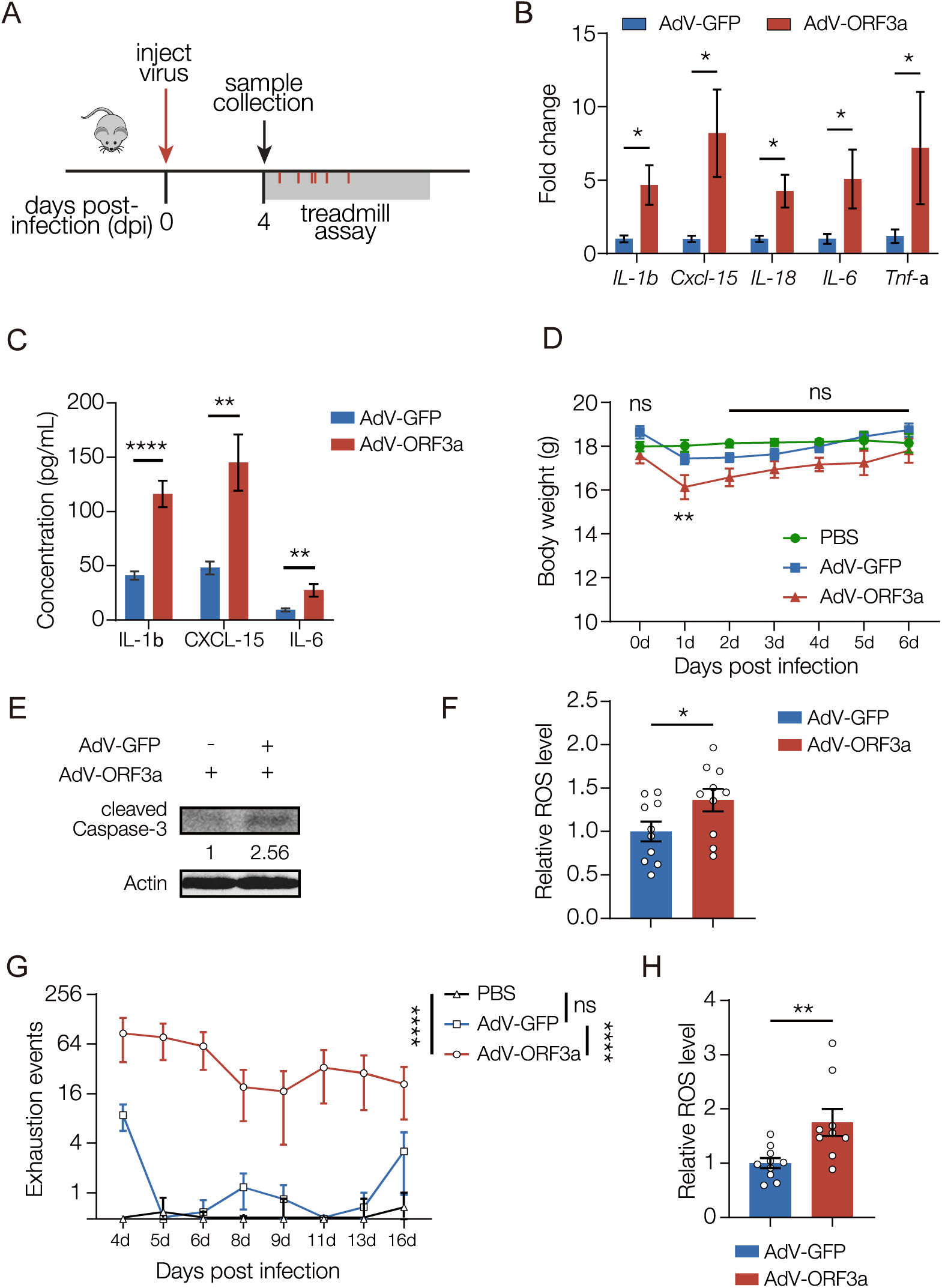
ORF3a induces neuroinflammation and reduces muscle performance in mice. **A.** Experimental design to study the skeletal muscle response to ORF3a expression in mice. **B.** qRT-PCR. Transcripts encoding cytokines were enriched in the brain of AdV-ORF3a.GFP infected mice compared to AdV-GFP infected controls. n≥5. **C.** Enzyme-linked immunosorbent assay (ELISA). Cytokine protein levels were enriched in in the brain of AdV-ORF3a.GFP infected mice compared to AdV-GFP infected controls. n≥5. **D.** Body weight changes over time. Mice infected with AdV-ORF3a.GFP showed a temporary drop in body weight at 1dpi compared to PBS injected or AdV-GFP injected controls. n=6. **E.** Western blot. GFP-positive brain lysates from AdV-ORF3a.GFP infected mice contained more cleaved Caspase-3 than AdV-GFP infected mice. **F.** ROS assay. H2DCFDA was used to measure ROS levels in GFP-positive brain tissue. AdV-ORF3a.GFP infected mice produced more ROS than AdV-GFP infected mice. Each data point represents an individual mouse. **G.** Longitudinal forced treadmill-running assay. AdV-ORF3a.GFP infected mice showed more exhaustion events than AdV-GFP infected mice or PBS injected mice. n≥11. **H.** ROS assay. H2DCFDA was used to measure ROS levels in skeletal muscle. AdV-ORF3a.GFP infected mice produced more ROS than AdV-GFP infected mice. Each data point represents an individual mouse. Significance was determined by two-sided unpaired Student’s t-test (B, C, F, H), and one-way ANOVA (D, G). Error bars represent SEM. (*) p< 0.05, (**) p< 0.01, (****) p < 0.0001, (ns) non-significant.

### A neurotoxic protein associated with Alzheimer’s disease activates the brain-muscle axis

Neuroinflammation can also be activated in the absence of infection. For example, Alzheimer’s disease (AD) is a non-infectious disease, and patients with AD often show chronic neuroinflammation, muscle weakness, and elevated ROS levels in the brain (Lai et al., 2017; Suryadevara et al., 2020). We performed a meta-analysis of 12 studies that characterized serum protein levels in a total of 585 AD patients and 439 healthy controls, and found AD patients had higher levels of IL-6 than unaffected patients (Fig. S6A). These observations suggested AD induces IL-6 expression and activates the brain-muscle signaling axis. Amyloid-β (Aβ42) is a neurotoxic protein involved in AD progression, and Aβ42 has been used to model AD in worms, flies, and mice. To understand if AD-associated neuroinflammation activates the brain-muscle axis, we expressed Aβ42 in the CNS of adult flies (hereafter AD flies). AD flies showed significantly reduced muscle performance increased ROS levels in the brain (Fig. 7A,B). Similar to ORF3a expressing flies, AD flies had reduced muscle mitochondrial activity but normal myofiber morphology (Fig.7C,D; Fig.S6B)(Casas-Tinto et al., 2011). In addition, AD flies showed elevated *upd3* expression in the brain, and increased *socs36E* expression in muscle (Fig. 7F). AD flies also showed enriched *10XSTAT92E.GFP* expression in muscle (Fig. 7G,H). These data argue Aβ42-induced neuroinflammation activates the brain-muscle signaling axis and disrupts muscle performance.

**Figure 7.**
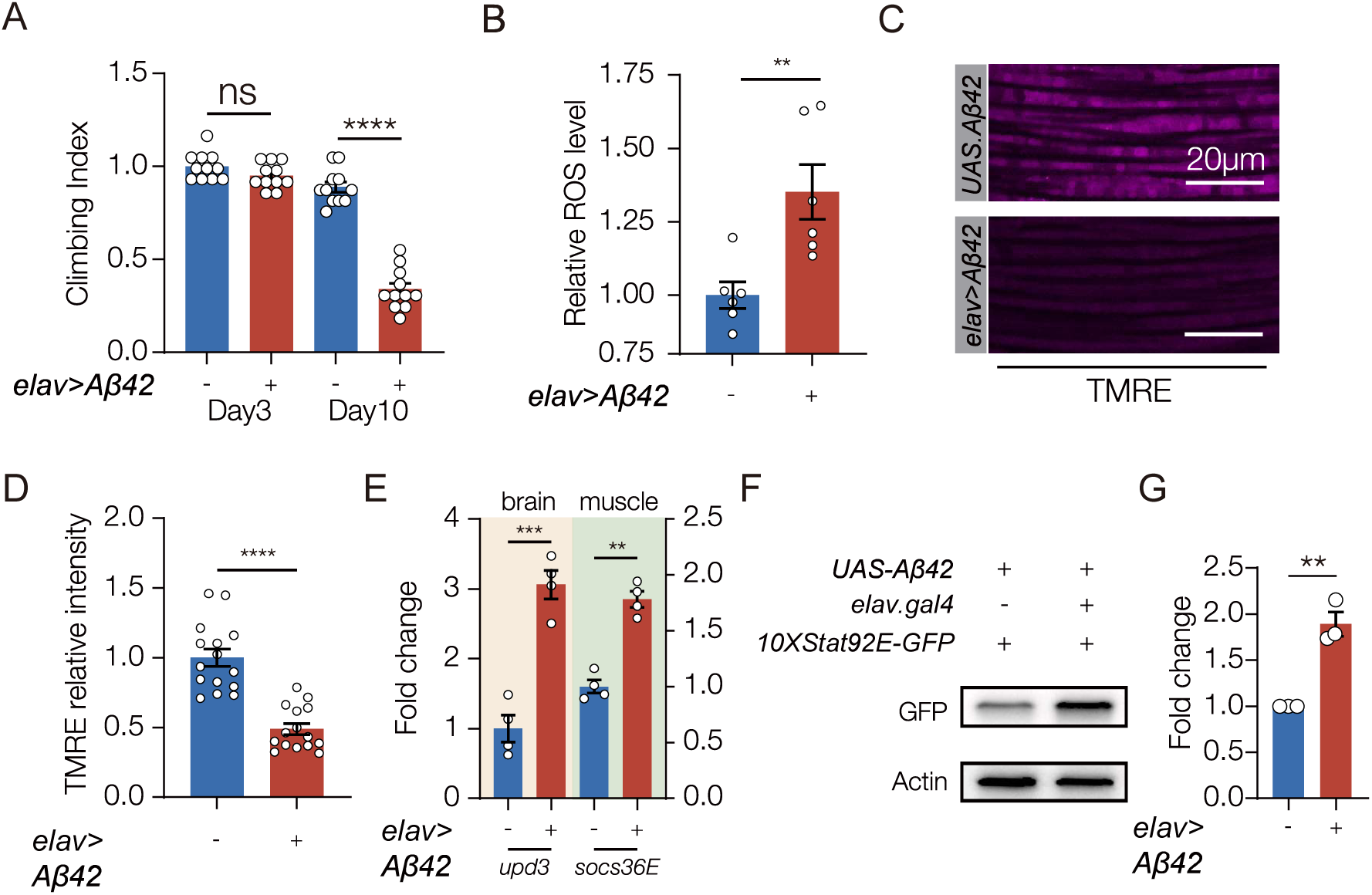
The brain-muscle axis is activated in a *Drosophila* model of Alzheimer’s disease (AD). **A.** Climbing index. Flies that expressed the AD-associated protein *Aβ 42* in the CNS (*elav>Aβ 42*) showed reduced climbing capacity compared to controls (*UAS.Aβ 42*) 10 days after eclosion. **B.** ROS assay. H2DCFDA was used to measure ROS in the brain. *elav> Aβ 42* flies produced more ROS than controls (*UAS.Aβ 42*). **C.** Confocal micrographs of indirect flight muscles stained with TMRE (violet) to assess mitochondrial membrane potential. Muscles from *elav>Aβ 42* flies showed less TMRE staining than controls (*UAS.Aβ 42*). **D.** Quantification of mitochondrial membrane potential shown in C. **E.** qRT-PCR. *elav>Aβ 42* flies expressed more *upd3* in the brain and more *socs36e* in indirect flight muscles than control flies (*UAS.Aβ 42*). **F.** Western blot. GFP expression from the JAK/Stat activity reporter *10XStat92E.GFP* in muscle was enriched in *elav>Aβ 42* flies compared to controls (*UAS.Aβ 42*). **G.** Quantification of GFP expression shown F. Experiments in B-G were performed on 10 day old flies. Significance was determined by one-way ANOVA (A), and two-sided unpaired Student’s t-test (B, D, F, H). For Western blots, qRT-PCR, and ROS assays relative expression was determined for a minimum of three biological replicates. See Fig. 1 legend for Climbing Index and TMRE data points. Error bars represent SEM. (**) p< 0.01, (***) p< 0.001, (****) p < 0.0001, (ns) non-significant.

## Discussion

We have shown that bacterial infections, the SARS-CoV-2 protein ORF3a, and the neurotoxic protein Aβ42 promote ROS accumulation in the brain. Excessive ROS induces Upd3 expression in insects, or IL-6 expression in mammals, and CNS-derived Upd3/IL-6 activates the JAK/Stat pathway in skeletal muscle. Muscle performance and mitochondrial activity are in turn reduced in response to JAK/Stat signaling. This is the first study to define a brain-muscle signaling axis, and highlights circulating Upd3/IL-6 as a novel mechanism by which the CNS communicates with skeletal muscle. Since infections and chronic diseases induce cytokine-dependent changes in muscle performance, our work suggests IL-6 could be a therapeutic target to treat muscle dysfunction caused by neuroinflammation.

Each of the models of neuroinflammation we developed or characterized has a clear clinical counterpart. For example, meningitis is an infection-induced inflammation of the meninges that is an important cause of mortality and morbidity in neonates and infants (Kim, 2010). The meninges act as a protective layer that surrounds the CNS, but gram-negative bacterial infections of the meninges, such as *E. coli*, can penetrate the blood-brain barrier and directly infect the CNS (Kim, 2008). Similar to our *Drosophila* model of bacterial-induced neuroinflammation, *E. coli* induced neonatal meningitis in rodents induced the expression of several cytokines, including the Upd3 orthologue IL-6 (Barichello et al., 2014), and inflammatory myopathies have been observed in patients recovering from meningeal infections (Martellosio et al., 2020). The Upd3/IL-6 brain-muscle signaling axis may therefore be activated in patients with CNS infections.

A common clinical symptom of COVID-19 is neuroinflammation, which can be assayed in the cerebrospinal fluid (CSF) of SARS-CoV-2 infected patients (Vanderheiden and Klein, 2022). There has been much debate as to whether COVID-19 related neuroinflammation is a systemic response to viral infections outside the CNS, or a more direct response to viral infection in the CNS (Crunfli et al., 2022; Song et al., 2021; Yang et al., 2021). A recent study in non-human primates argues SARS-CoV-2 can infect neurons (Beckman et al.), which supports our findings that ORF3a was expressed in the brain of patients with COVID-19. While our study does not reveal how SARS-CoV-2 proteins arrive in the CNS, our results argue ORF3a is neuroinvasive and causes neuroinflammation in flies and mammals. The ORF3a transgenic fly is thus a preclinical model of COVID-19 that does not require the biosafety level precautions needed for working with the SARS-CoV-2 virus.

Mild SARS-CoV-2 infection in the respiratory system was recently shown to promote neuroinflammation in the CSF, to enhance microglial reactivity, and to induce the loss of oligodendrocytes and myelinated axons in mice (Fernández-Castañeda et al., 2022). SARS-CoV-2 also induced changes in brain structure, visible by longitudinal magnetic resonance imaging in infected patients, and enhanced cognitive decline (Douaud et al., 2022). Interestingly, mild respiratory SARS-CoV-2 infection caused IL-6 levels to be enriched in the serum for several weeks post-infection (Fernández-Castañeda et al., 2022). It is possible that the brain-muscle axis is activated by respiratory infections via the CSF, and continues to signal long after the initial infection. Similar to respiratory infections, ORF3a expression in the brain activated neuroinflammation and reduced cell survivability in the CNS, suggesting neurodegeneration is a common pathology of COVID-19 regardless of the site of SARS-CoV-2 infection or viral protein expression.

During the post-acute phase of COVID-19, a significant proportion of patients experience symptoms outside the respiratory system, and this constellation of post-acute sequelae has been termed Long-COVID (Ballering et al., 2022; Subramanian et al., 2022; Xu et al., 2022) (Huang et al., 2021; Taquet et al., 2021). Cohort studies of COVID-19 patients documenting non-respiratory symptoms from three to twelve months after acute infection identified an increased incidence of neurologic disorders and somatic symptoms that include muscle pain and fatigue (Ballering et al., 2022; Xu et al., 2022). SARS-CoV-2 proteins, such as Spike protein, have been detected in circulation after the viral infection is cleared (Swank et al., 2022). We found ORF3a is neuroinvasive and induces neurodegeneration and reduced muscle function. Long-COVID symptoms could be caused by the persistent expression of SARS-CoV-2 proteins, including ORF3a, after the acute phase of SARS-CoV-2 infection has been cleared. By using ORF3a to model Long-COVID, we have found that SARS-CoV-2 proteins can have local effects on the infected tissue and far reaching effects on whole-body physiology. One underlying cause of Long-COVID could be the disruption of inter-organ communication pathways by the long-term expression of SARS-CoV-2 proteins during the post-acute phase of COVID-19. The Upd3/IL-6 brain-muscle signaling axis may therefore be activated by SARS-CoV-2 during both the acute and post-acute phases of COVID-19.

Alzheimer’s disease (AD) is a chronic neurodegenerative disorder that affects nearly six million people in the United States, and neuroinflammation is one factor that leads to AD pathology (Dhapola et al., 2021). AD also accelerates sarcopenia, which is the loss of skeletal muscle mass and function with aging (Beeri et al., 2021). Although the connections between AD and sarcopenia are poorly understood, AD and musculoskeletal aging are associated with an overlapping set of differentially expressed genes that regulate mitochondrial function (Giannos et al., 2022). In addition, skeletal muscle mitochondria from individuals in the early phases of AD showed reduced respiratory kinetics and poor control of mitochondria coupling (Morris et al., 2021). Chronic neuroinflammation in AD correlates with reduced mitochondrial function in skeletal muscle, and our in silico analysis argues circulating IL-6 levels are increased in AD patients. Upd3 expression was also enriched in the fly model of AD, suggesting the Upd3/IL-6 brain-muscle signaling axis may be active in AD patients

Our three disease models have identified IL-6 and Stat inhibitors as potential therapeutics for patients with CNS infections, COVID-19, and AD. In fact, we successfully mitigated the effects of CNS bacterial infections by inhibiting Upd3/IL-6 and the JAK/Stat pathway. The Stat inhibitor ruxolitinib is approved to treat alopecia, psoriasis, lymphoma, and myelofibrosis (Haq and Adnan, 2022). Myelofibrosis is defined a myeloproliferative neoplasm that is often correlated with variants in *JAK2* (Arber et al., 2016). Although musculoskeletal symptoms are not part of the myelofibrosis diagnostic criteria, myelofibrosis patients that received ruxolitinib showed significant improvement of muscle mass over a six-month longitudinal study (Lucijanic et al., 2021). These clinical results align with our overall model that IL-6 and Stat inhibitors could inhibit changes to muscle performance induced by the Upd3/IL-6 brain-muscle signaling axis, and prevent muscle atrophy associated with muscle disuse.

### Evolution of the brain-muscle axis

Why then did the Upd3/IL-6 brain-muscle signaling axis evolve? Skeletal muscle comprises 45% of the total human body mass, and consumes 28% of the body’s total energy at rest (Chen et al., 2021b; McClave and Snider, 2001b). The adult brain consumes 20% of the resting energy expenditure (McClave and Snider, 2001a). The brain-muscle axis might limit mitochondrial function in muscle to reduce energy consumption and redirect energy expenditure toward the CNS for recovery after disease or injury-induced neuroinflammation. The immune system also consumes extensive energy, and shifts from 11% total energy consumption in a healthy adult to 43% energy consumption during infection (Straub, 2017). The brain-muscle signaling axis may induce muscle weakness to redirect energy expenditure toward the immune system to accelerate clearance of infections or to promote other immune system functions necessary for recovery.

A second hypothesis is that the brain-muscle axis restricts muscle function to protect skeletal muscle from contraction-induced injuries. Muscle strains are contraction-induced injuries, and are the most common sports injury (Dueweke et al., 2017). While neuroinflammation has not been associated with contraction-induced injuries, it is possible that CNS infections and chronic diseases inadvertently cause hypercontracility and muscle strains. If the brain-muscle axis reduces mitochondrial function in response to neuroinflammation, then skeletal muscle would be protected from contraction-induced injury. Since four drugs targeting neuroinflammation are currently in clinical trials (Dhapola et al., 2021), it will important to identify the positive effects of the brain-muscle signaling axis on CNS recovery, systemic immunity, and the prevention of muscle injuries.

## Materials and methods

### Drosophila genetics

The wild-type flies used were *w^1118^* (Bloomington Stock Center, 3605) unless specifically mentioned, all flies were maintained on standard cornmeal fly food unless noted otherwise. Flies were cultured at 25℃under a normal light/dark cycle, unless noted otherwise. The *Drosophila* stocks used in this study are described in Flybase (http://flybase.org/) unless specified.

The RNAi lines were *UAS-Upd3-RNAi* (Bloomington Stock Center, 32859) and *UAS-Dome-RNAi* (Bloomington Stock Center, 34618). Gal4 line were *GS.elav-Gal4* (Bloomington Stock Center, 43642), *GS.nSyb-Gal4* (Bloomington Stock Center, 80699), *elav-Gal4,Sb/TM6b* (Dr. Helen McNeill, WUSM), *elav^strong^-Gal4* (Bloomington Stock Center, 458), *elav-Gal4,UAS-sfCherry/Cyo;TM3/TM6b* (Bloomington Stock Center, 93287), *repo-Gal4* (Bloomington Stock Center, 7415), and *Mef2-Gal4* (Bloomington Stock Center, 50742). UAS lines were *UAS-ORF3a* (lab made), *UAS-Upd3* (Dr. Bruce Edgar, The University of Utah), *UAS-PGRP-LCa* (Bloomington Stock Center, 30917), *UAS-PGRP-LE* (Bloomington Stock Center, 33054), *UAS-p35* (Bloomington Stock Center, 5072), UAS-Abeta42 (PMID: 21389082). Mutant lines were *imd^1^* (Bloomington Stock Center, 55711), *dorsal^1^* (Bloomington Stock Center, 3236), *dif^1^* (Bloomington Stock Center, 36559). *10XStat92E-GFP* reporter line was used to analyze JAK/Stat pathway activation (Bloomington Stock Center, 26197).

UAS-SARS-CoV-2-ORF3a transgenic flies were generated by PCR-mediated subcloning of the SARS-CoV-2-ORF3a coding sequence (pDONR207-SARS-CoV-2 ORF3a, #141271, Addgene) into pUASt-Attb (EcoRI/XbaI). ORF3a was amplified with Takara PrimerSTAR PCR enzyme (R050B, Takara) using the following primers:

ORF3a-CDS-Forward-CGGAATTCATGGACCTGTTCATGAGAATCTT

ORF3a-CDS-Rerverse-GCTCTAGATTACAGTGGCACGGAGGTG

Plasmid DNA was injected and targeted to a C31 integration site (Bloomington Stock 24481, Rainbow Transgenic Flies); stable insertions were identified by standard methods.

### Mice

Mice protocols used in this study were approved by the Institutional Animal Care and Use Committee of Tsinghua University and performed in accordance with their guidelines (approval number:22-CG1). The laboratory animal facility has been accredited by the Association for Assessment and Accreditation of Laboratory Animal Care International. Six-week-old C57BL/6 mice were purchased from Charles River. All animals were maintained in a specific pathogen-free animal facility at Tsinghua University with a 12h light/dark cycle and normal chow diet and water.

### Longevity and motor function assays

1d old adult flies were collected and transferred to fresh food daily for both assays. For longevity analysis, the number of dead flies was recorded daily. Kaplan-Meier survival curves were generated, and statistical analysis was performed using log-rank analysis (Prism9, GraphPad Software). To assess motor function, climbing assays were performed as described(Moore et al., 2018). Briefly, 15-20 flies were placed into empty vials (9.5 cm high,1.7 cm in diameter) with flat bottoms, the flies were forced to the bottom of a vial by firmly tapping the vial against the bench surface. Eight seconds after the tap, the number of flies that climbed up the walls of a vial above the 5-cm mark was recorded as positive.

To assess climbing speed, around 20 adult flies were transferred to a 100ml glass cylinder. The cylinder was tapped rapidly for 4 times. After the final tap, fly movement was recorded by a camera. Average climbing speed was calculated by speed=height/8s.

For mice treadmill assay, 6-week-old mice were familiarized with the treadmill environment before experiment, and assayed by a six-channel motor-drive uphill treadmill (UGO Basile, Varese, Italy) at the speed of 15 m/min for 5 min/day throughout the experiment. The grid (3mm bars, placed 8mm apart.) attached to mouse assembly, delivers the light foot-shock (1mA). Shock intensity and frequency were recorded.

### Circadian rhythm analysis

To analyze the sleep patterns of flies expressing either ORF3a or a control LacZ transgene in neuronal or glial cells, we used the *Drosophila* activity monitor (DAM) system (Trikinetics, Waltham). For this, we counted the number of times flies cross infrared beams in a 7-day period. Individual flies were gently inserted in capillary tubes (5mm) containing 5% sucrose and 2% agar. These tubes were then loaded horizontally in DM2 Trikinetics monitors under 12-hour Light/12-hour Dark conditions at 25°C. Counts were recorded by DAMSystem 3.11.1 and subsequently scanned and binned into 1-min intervals using DAMFileScan 1.11. Actograms were plotted using NIH’s ImageJ plugin ActogramJ (https://bene51.github.io/ActogramJ/). Total activities of at least 6 consecutive days were plotted.

### Bacteria, virus, and infection

Collect *E.coli* (LB medium, 37°C) at optimal status (OD_600_ = 0.6∼0.8) by centrifuged at 6000 g for 5min at room temperature, wash the bacteria with sterile PBS for three times, and then dilute the bacteria pellet with PBS to OD_600_ = 200. Infection was performed as described by using 0.0125 mm needles (Fine Science Tools, 2600210)(Cao et al., 2013). Flies were kept at 29°C and flipped daily with standard fly food.

AdV-ORF3a was construed in Keda Biotech, 20ul 3x10^10^PFU/ml virus was delivered to wildtype mice (C57BL/6) by retro-orbital injection.

### Reactive oxygen species detection

H2DCFDA (Thermo Fisher, D399) was used for ROS detection. For total ROS, 5 fly heads were ground in 100ul cold PBS, then centrifuged for 15min (13500g, 4 degrees). 50ul supernatant was incubated with 150ul H2DCFDA working solution (14uM, in PBS) for 30 min at 37 degrees. Fluorescence was measured by BioTek Synergy H1 (Ex/Em: 488/525).

### N-acetyl-L-cysteine treatment

1g N-acetyl-L-cysteine (NAC) was dissolved in 10mL water, the solution could be aliquoted into 1 ml per EP tube and frozen or stored at -80 degrees. For larvae treatment, 4g common fly food was mixed with 200ul 1x NAC solution, and incubated at 25 degrees for 2∼2 days egg laying. For adult treatment, 100ul of 0.5xNAC solution was dipped onto the surface of fresh food in vials (diameter: 2 cm). The vials were then allowed to dry at room temperature for 6 hours. Female adult flies were cultured on this NAC food at 25 degree, and fresh food was changed every day.

### Mifepristone (RU486) treatment

In RU486-induced experiments, 20 adult female flies (2–4 days old) per vial were fed with 50ug/mL of RU486 (M8046, Sigma; stock solution is 10mg/mL in DMSO, dilute to working solution with ethanol before use) for 6 days. 100ul working solution was added to the surface of fresh food (2cm diameter) and evaporate overnight (room temperature, protect from light).

Flies were raised on RU486/buffer-contained food at 25 degrees, and fresh food was changed every day.

### Plasmids

pCMV-GFP-SARS-CoV-ORF3a was generated by recombining the SARS-CoV-2-ORF3a coding sequence (pDONR207 SARS-CoV-2 ORF3aA, #141271, Addgene) into pDEST-CMV-N-EGFP (#122842, Addgene). pCMV-EGFP-ORF3a-Q57E-S58L-Q116L was constructed as described(Yang et al., 2022). Following primers were used: Q57E-F: CTGCTGGCCGTGTTCGAGTCCGCCTCTAAGATC

Q57E-R:GATCTTAGAGGCGGACTCGAACACGGCCAGCAG

Q57E-S58L-F: CTGGCCGTGTTCGAGCTCGCCTCTAAGATCATC

Q57E-S58L-R: GATGATCTTAGAGGCGAGCTCGAACACGGCCAG

S116L-F: CTCTGGTGTACTTCCTGCTGAGCATCAACTTCGTGCG

S116L-R: CGCACGAAGTTGATGCTCAGCAGGAAGTACACCAGAG

### LysoTracker staining

Hela cells were seeded in 6-well plates with cover slips and grown to 50% confluency at 37°C with 5% CO_2_ in Dulbecco’s modified Eagle’s medium (12430047, Invitrogen) supplemented with 10% heat-inactivated FBS (A4766801, Invitrogen). Cells were then transfected with 1000ng DNA, using standard Lipofectamine 3000 protocol (L3000008, Invitrogen). 24h post transfection, media was removed, and cells were incubated with 10 nM LysoTracker Red DND-99 (L7528, Invitrogen) for 1h. Cells was washed with PBS for three times, mounted and imaged with a Zeiss LSM800 confocal microscope.

### TMRE and phalloidin staining

For TMRE staining of muscle, adult fly hemi-thoraces were dissected in S2 medium (without serum), and incubated in TMRE staining solution (200 nm TMRE in S2 medium, T669, Thermo Fisher Scientific) for 30 min at room temperature. For TMRE staining of cells, Hela cells were seeded into 6-well-plate with a cover slip, and transfected with 1000ng pCMV-GFP-ORF3a or pCMV-GFP-ORF3a-Q57H-S58L-Q116L for 24h. Cells were washed with PBS and incubated with TMRE staining solution (50 nm TMRE in DMEM, without serum) for 15 min at 37 degree. After staining, samples (muscle or cells) were washed with PBS for 5 times, and immediately imaged using a Zeiss LSM880 confocal microscope. TMRE intensity was quantified using ImageJ software and normalized to corresponding control group. For phalloidin staining, adult fly muscles were dissected in PBS, and fixed with 4% formaldehyde for 12min, and tissue staining was performed as described (Yang et al., 2022). Alexa Fluor™ 555 Phalloidin (1:40; A34055, Thermo Fisher Scientific) was used to visualize F-Actin. Tissues were imaged with a Zeiss LSM800 confocal microscope.

### Immunohistochemistry

For patient samples, single-blind immunohistochemical was performed by Wuhan Servicebio technology with standard method as described (Watson and Soilleux, 2015), except anti-SARS-CoV-2 ORF3a antibody was used (1:500; PA5-116946; Thermo Fisher Scientific).

### Western blotting

To detect ORF3a and c-Caspase-3 expression, 10 adult female heads were homogenized in 200 ul IP buffer (20 mM Hepes, pH=7.4, 150 nM NaCl, 1% NP40, 1.5 mM MgCl2, 2 mM EGTA, 10 mM NaF, 1 mM Na3VO4, 1X proteinase inhibitor). To detect 10XSTAT-GFP reporter expression, 8 adult thoraces (4 male and 4 female) were homogenized in 200ul IP buffer. Samples were incubated on ice for 30 min and large debris was removed by 15min centrifugation (12,000xg). Anti-cleaved-Caspase 3 (1:500; #9661, Cell Signaling Technology), anti-SARS-CoV-2-ORF3a (1:250; 101AP, FabGennix International Inc), anti-Actin (1:500; JLA20, DSHB), anti-beta-Tubulin (1:500; E7-C, DSHB), and anti-GFP (1:1000; TP-401, Torrey Pines Biolabs) were used for immunoblotting. Western blots were performed by standard method using precast gels (#456-1096, BioRad), and imaged with the ChemiDoc XRS+ system (BioRad).

### Cytokine protein measurements

Samples were prepared with standard method as described (Sukoff Rizzo et al., 2012). Total protein was determined by BCA Protein Assay Kit (23225, Thermo Scientific). Cytokines and chemokines were analyzed for by Elisa (IL-1beta, ab197742, Abcam; CXCL-15, MOFI01258, Assay Genie; IL-6, A12219, Yojanbio).

### Quantitative real-time RT-PCR

Total RNA was extracted with TRIzol (15596026, Invitrogen), and quantitated with a Nanodrop 2000 (Thermo Fisher). cDNA was prepared by reverse transcription with All-in-One 5X RT MasterMix (G592, Applied Biological Materials Inc) with 1000ng RNA. BlasTaq 2X qPCR MasterMix (G891, Applied Biological Materials Inc) and ABI Step One system (Applied Biosystems) were used for quantitative RT-PCR. Quantification was normalized to endogenous ribosomal protein Rp32 mRNA or GAPDH mRNA. RT-PCR primers included:

Diptercin-F: GGCTTATCCGATGCCCGACG

Diptercin-R: TCTGTAGGTGTAGGTGCTTCCC

Attacin-A-F: ACGCCCGGAGTGAAGGATGTT

Attacin-A-R: GGGCGATGACCAGAGATTAGCAC

Drosomycin-F: GCAGATCAAGTACTTGTTCGCCC

Drosomycin-R: CTTCGCACCAGCACTTCAGACTGG

Metchnikowin-F: GACGCAACTTAATCTTGGAGCG

Metchnikowin-R: TTAATAAATTGGACCCGGTCTTGGTTGG

Eiger-F: AGCGGCGTATTGAGCTGGAG

Eiger-R: TCGTCGTCCGAGCTGTCAAC

4E-BP-F: GTTTGGTGCCTCCAGGAGTGG

4E-BP-R: CGTCCAGCGGAAAGTTTTCG

Mmp2-F: GAGATGCCCATTTCGATGCG

Mmp2-R: GCCGTACAACTGCTGAATGC

Duox-F: GCCCTGCTGCTTCTACTGAT

Duox-R: CGCTGTTTCTCGGTCTGACT

Nox-F: TCCGCAAGCTATTCCTGGAC

Nox-R: TTGCTCGGCAAAGTCCATCT

Sod1-F: TGCGTAATTAACGGCGATGC

Sod1-R: CATGCTCCTTGCCATACGGA

Sod2-F: CAAGTCGAAGAGCGACACCA

Sod2-R: TTGTTGGGCGAGAGGTTCTG

Cat-F: CAAAATGGCTGGACGCGATG

Cat-R: GGGAGGCATCCTTGATTCCA

Upd3-F: CCTGCCCCGTCTGAATCTCA

Upd3-R: TGAAGGCGCCCACGTAAATC

Socs36E-F: AGGAGGAGTTCCTCTTCTCGGTC

Socs36E-R: CGTGGCAGTCGAAGCTGAAC

CXCL-15-F: TCCAGAGCTTGAAGGTGTTGCC

CXCL-15-R: AACCAAGGGAGCTTCAGGGTCA

IL-6-F: CACAAGTCCGGAGAGGAGAC

IL-6-R: CAGAATTGCCATTGCACAAC

IL-18-F: GACCAAGTTCTCTTCGTTGACAA

IL-18-R: ACAGCCAGTCCTCTTACTTCA

IL-1beta-F: CGCAGCAGCACATCAACAAGAGC

IL-1beta-R: TGTCCTCATCCTGGAAGGTCCACG

TNF-F: CGTGGAACTGGCAGAAGAG

TNF-R: TGAGAAGAGGCTGAGACATAGG

ORF3a-F: CGCTACTGCAACGATACCGA

ORF3a-R: AACAGCAAGAAGTGCAACGC

### Ethics statement

Research participants were enrolled at Wuhan Jinyintan Hospital of Huazhong University of Science and Technology (HUST) through Human Investigation Committee Protocols, KY-2020-15.0. The Institutional Review Board at HUST approved the protocols, and informed consent was obtained from all participants. Postmortem COVID-19 brain tissues were obtained from Department of Forensic Medicine Biobank of Tongji Medical college in HUST. This study was supported by the Emergency Novel Coronavirus Pneumonia Project from the Ministry of Science and Technology of China (2020YFC0844700).

### Bioinformatic and statistical analysis

Protein structure was generated by Chimera1.16 (USCF). The AD meta analysis and Forest Plot was generated with RevMan5 software. Comparisons of two samples were made using Student’s t-test, and multiple samples by one-way or two-way ANOVA. Survival curves were compared using the Kaplan–Meier test. P values of less than 0.05 were considered statistically significant. All statistical analyses were performed with GraphPad Prism 9 software. The sample sizes and number of replicates are indicated in the figure legends. Data collection and data analysis were routinely performed by different authors to prevent potential bias. All individuals were included in data analysis.

## Acknowledgments

We thank Dr. Bruce Edgar (The University of Utah) for the kind gift of *UAS-Upd3* stock; Dr. Helen McNeill (WUSM) for providing stock of *elav-Gal4,Sb/TM6b*. We also thank Helen McNeill and Mayssa Mokalled for critical reading of the manuscript. This work was funded by grants from the National Institutes of Health (R01AR070299) to A.N.J.; the National Natural Science Foundation of China (32188101, 81961160737, 31825001, 81730063, 8191101056, 82041006 and 31700148) to G.C.; the National Key Research and Development Plan of China (2021YFC2300200, 2020YFC1200104, 2018YFA0507202) to G.C.; Tsinghua University Spring Breeze Fund (2020Z99CFG017), Shenzhen Science and Technology Project (JSGG20191129144225464) to G.C.; Innovation Team Project of Yunnan Science and Technology Department (202105AE160020) and the Yunnan Cheng Gong expert workstation (202005AF150034) to G.C.; Natural Science Foundation of Heilongjiang Province (JQ2021C005) to X.Y.; the National Institutes of Health (R01AG059871) to D.R..

## Author Contributions

Conceptualization, S.Y., G.C., and A.J.; Methodology and Validation, S.Y., M.T, Y.D., D.C., D.R., and A.J.; Formal Analysis, S.Y., M.T, Y.D., S.F., Y.W., D.C., T.O., J.M., W.L., Z.Y., D.R., X.Y., W.T., G.C. and A.J.; Virtualization, S.Y., Z.Y., G.C., and A.J.; Investigation, S.Y., M.T, Y.D., S.F., Y.W., and D.C.; Resources, D.R., X.Y., W.T., G.C. and A.J.; Writing Original Draft, S.Y., X.Y., W.T., G.C. and A.J.; Review & Editing, S.Y., M.T, Y.D., S.F., Y.W., D.C., T.O., J.M., W.L., Z.Y., D.R., X.Y., W.T., G.C. and A.J.; Supervision, S.Y., X.Y., W.T., G.C. and A.J.; Funding Acquisition, D.R., X.Y., G.C. and A.J..

## Competing interests

The authors declare no competing interests.

## Data availability

The data that support the findings of this study are available from the corresponding authors upon request.

---

**Figure S1. Related to Figure 1.**
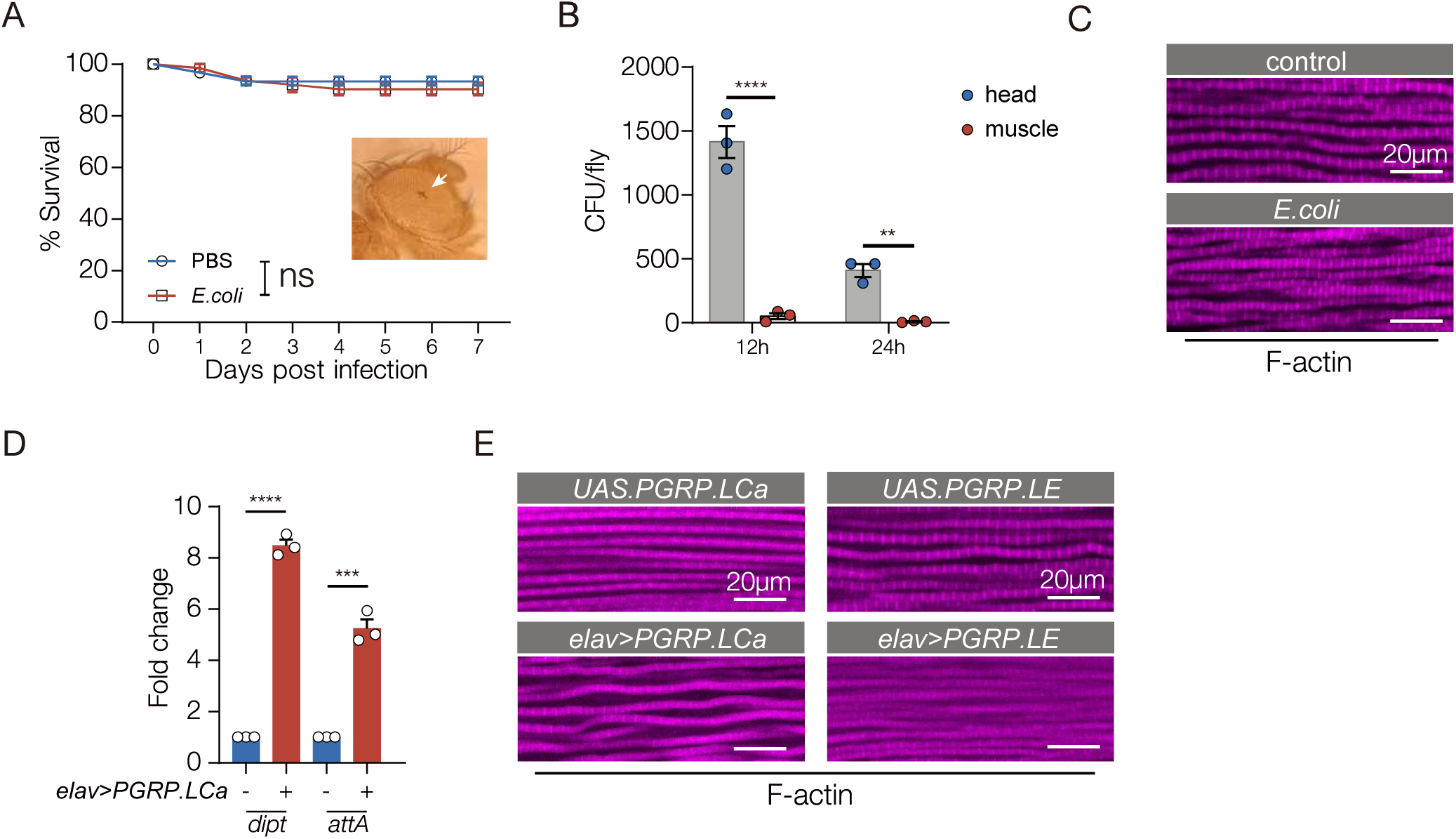
**A.** Survival curve. Sterile-injected (PBS, control) and *E. coli* injected flies showed comparable longevity. White arrow shows the injection site. n=3 cohorts per genotype, with n≥20 flies per cohort. **B.** Colony forming unit (CFU) assay. Bacterial load in the brain dramatically decreased between 12- and 24-hours after *E. coli* injection. Muscle showed minimal bacterial infection. Each data point represents one biological replicate, with n=5 flies per replicate. **C.** Confocal micrographs of indirect flight muscle stained with phalloidin to visualize F-actin (violet). Muscle morphology was comparable between sterile-injected (PBS, control) and *E. coli* injected flies. **D.** qRT-PCR. Transcripts encoding the antimicrobial peptides *Dipt* and *attA* in the brain was enriched in *elav>PGRPLCa* flies compared to controls (*UAS.PGRPLCa*). Data points represent independent biological replicates, with n≥10 flies per cohort. **E.** Confocal micrographs of indirect flight muscle stained with phalloidin to visualize F-actin (violet). Muscle morphology was comparable among *elav>PGRPLCa* flies, *elav>PGRPLE* flies, and controls (*UAS.PGRPLCa, UAS.PGRPLE*). Significance was determined by Kaplan–Meier tests (A), or two-way ANOVA (B, D). Error bars represent SEM. (*) p< 0.05, (**) p< 0.01, (***) p< 0.001, (****) p < 0.0001, (ns) non-significant.

**Figure S2. Related to Figure 2.**
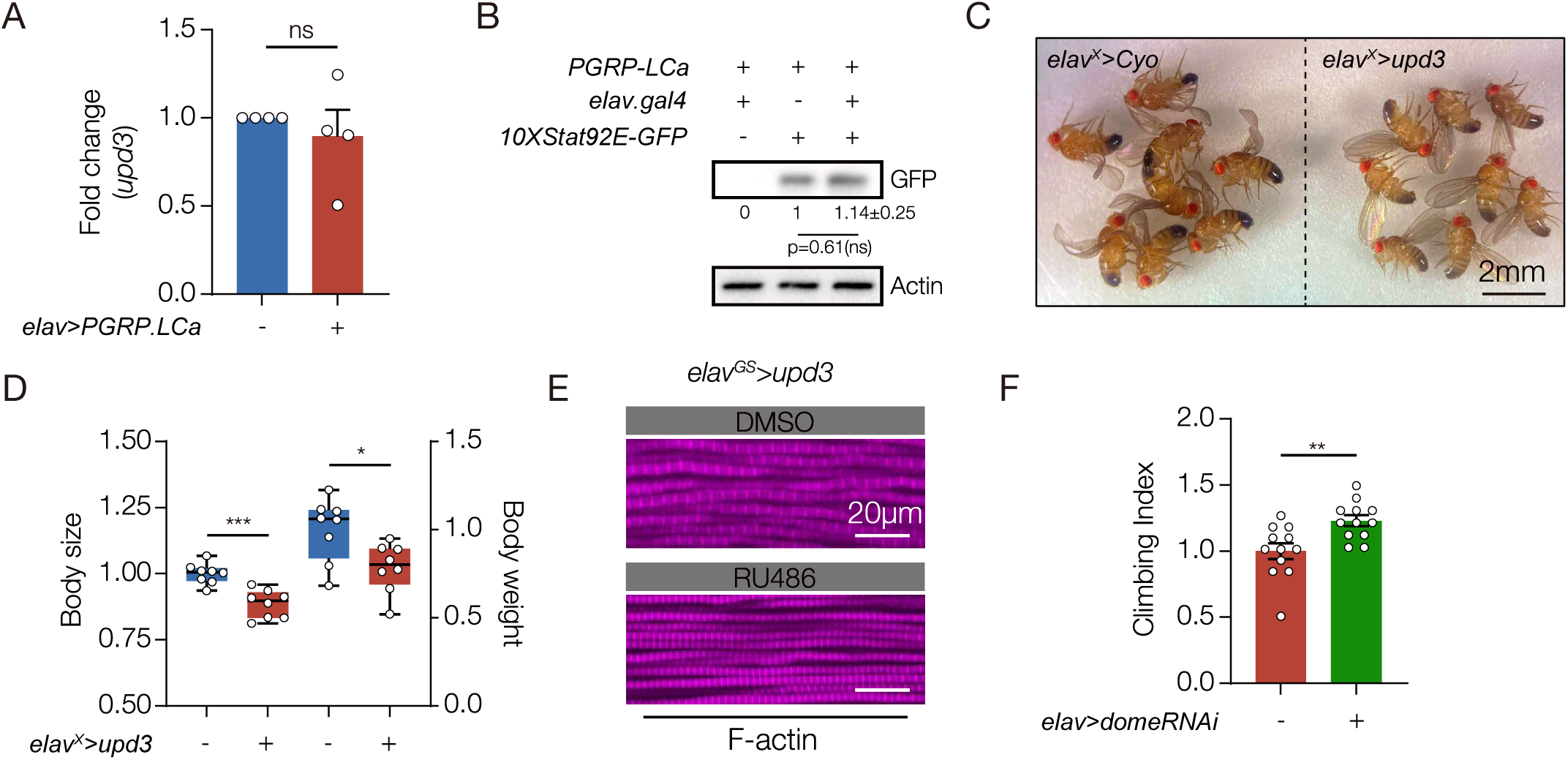
**A.** qRT-PCR. Flies that expressed PGRP in the CNS (*elav>PGRP.LCa)* showed similar levels of *upd3* mRNA in the brain as control flies. **B.** Western blot. GFP expression from the JAK/Stat activity reporter *10XStat92E.GFP* in muscle was similar between control and *elav>PGRP.LCa* flies. Relative expression was determined for three biological replicates. **C.** Micrographs of adult flies 3-5 days after eclosion. *elav^X^>upd3* flies were smaller than controls. **D.** Normalized body size (left Y axis) and body weight (right Y axis) of control and *elav^X^>upd3* flies. **E.** Confocal micrographs of indirect flight muscles stained with phalloidin to visualize F-actin (violet). *elav^GS^>upd3* flies treated with RU486 or DMSO showed similar myofiber morphology **F.** Climbing index. *dome^RNAi^* was used to knock down Dome expression in skeletal muscle of *E. coli* infected flies. Infected flies with reduced Dome expression (*Mef2>dome ^RNAi^*) showed improved climbing capacity compared to controls at 2 dpi. Significance was determined by two-sided unpaired student’s t-test. For qRT-PCR, data points represent biological replicates, with n≥10 flies per cohort. See Fig. 1 legend for Climbing Index data points. Error bars represent SEM. (*) p< 0.05, (**) p< 0.01, (***) p< 0.001, (****) p < 0.0001, (ns) non-significant.

**Figure S3 Related to Figure 4.**
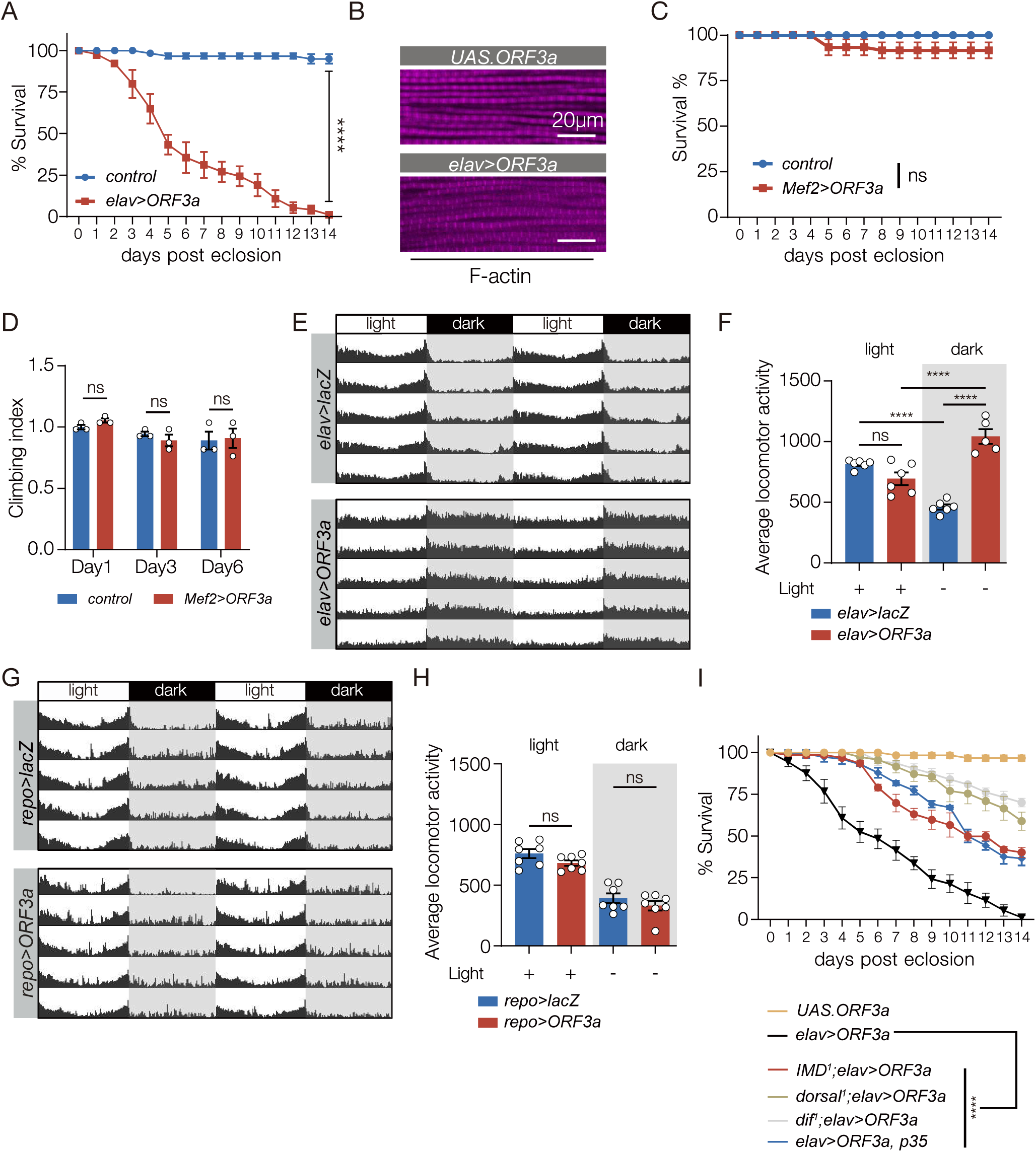
**A.** Survival curves. *elav>ORF3a* flies showed a significant reduction in longevity compared to control flies (*UAS.ORF3a*). n=5 cohorts per genotype, with n≥20 flies per cohort. **B.** Confocal micrographs of indirect flight muscles stained with phalloidin to visualize F-actin (violet). *elav>ORF3a* flies and control flies (*UAS.ORF3a*) showed similar myofiber morphology. **C.** Survival curve. *Mef2>ORF3a* flies and control flies (*UAS.ORF3a*) showed comparable longevity. n=3 cohorts per genotype, with n≥20 flies per cohort. **D.** Climbing index. *Mef2>ORF3a* flies showed and control flies (*UAS.ORF3a*) showed similar climbing capacity at 1-, 3-, and 6-days after eclosion. **E,G.** Actograms. Average activity of flies over 2-days is shown. **E.** Flies that expressed ORF3a broadly in the CNS (*elav>ORF3a*) flies were more active in dark cycles than control flies (*elav>lacZ*). **F.** Quantification of data shown in E. n = 28 flies per each genotype. **G.** Flies that expressed ORF3a only in glial cells (*repo>ORF3a*) flies showed similar activity in light and dark cycles as control flies (*repo>lacZ*). **H.** Quantification of data shown in G. n = 16 flies per each genotype. **I.** Survival curves. *elav>ORF3a* flies with homozygous mutations affecting the IMD pathway (*IMD^1^*) or the Toll pathway (*dorsal^1^*, *dif^1^*) showed improved longevity compared to *elav>ORF3a* flies. *elav>ORF3a* flies that expressed the inhibitor of apoptosis p35 in the CNS also showed improved longevity compared to *elav>ORF3a* flies. n=3 cohorts per genotype, with n≥20 flies per cohort. Significance was determined by Kaplan–Meier test (A, C, I), and two-way ANOVA (D, F, H). Data represent the average of at least three independent tests. Error bars represent SEM. (****) p < 0.0001, (ns) not significant.

**Figure S4 Related to Figure 5.**
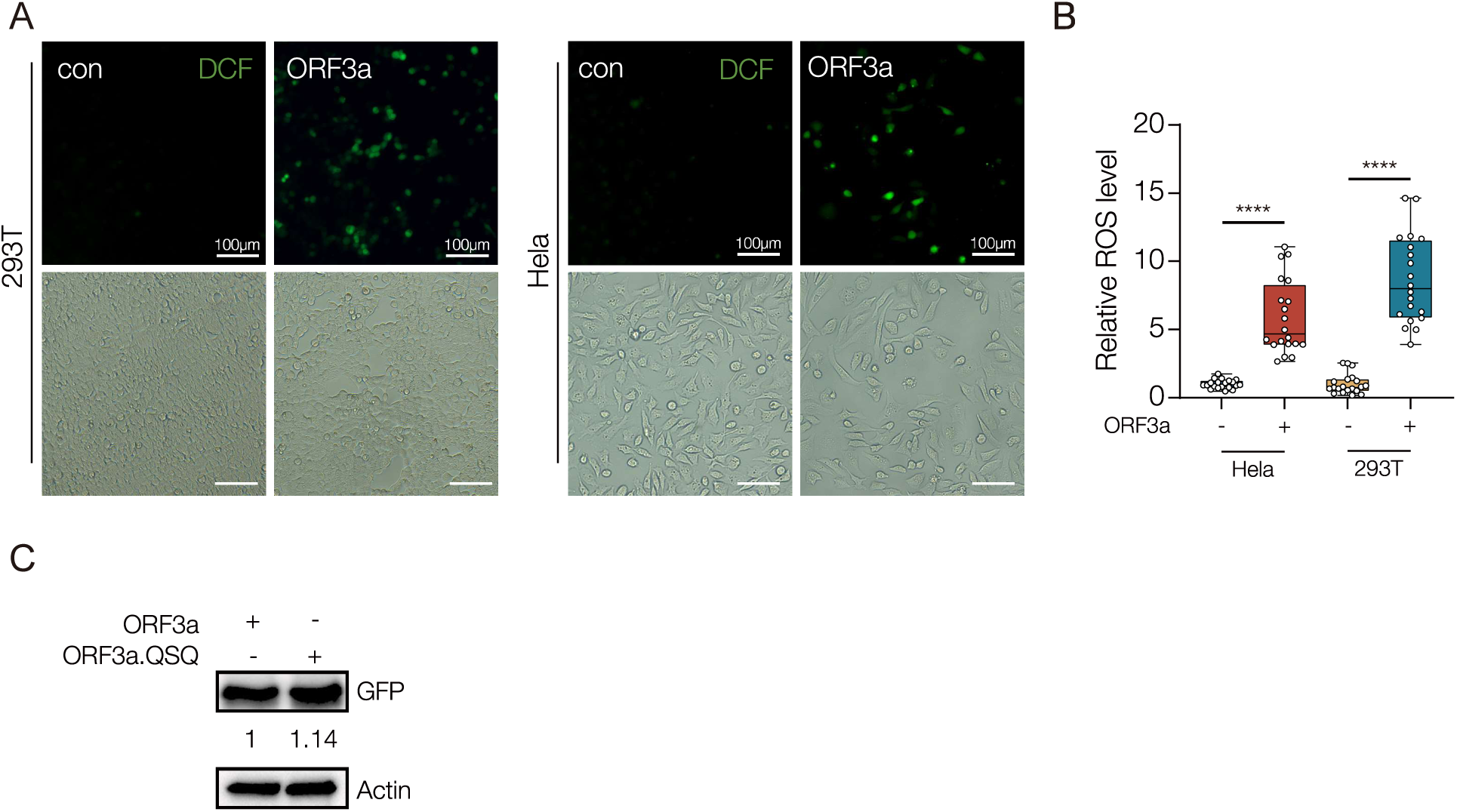
**A.** ROS assay. H2DCFDA was used to measure ROS in cultured cells. Micrographs of DCF fluorescence (green) in HEK293T cells (left) and HeLa cells (right) transfected with wild-type ORF3a. ORF3a transfected cells produced more ROS than untransfected controls. **B.** Quantification of ROS levels shown in A. Data points represent fluorescence in a single cell normalized to control cells. n=5 fields **C.** Western blot. HEK293T cells transfected with wild-type ORF3a and ORF3a.QSQ showed similar levels of ORF3a protein expression. Significance was determined by two-sided unpaired student’s t-test (B). Error bars represent SEM. (****) p< 0.0001.

**Figure S5 Related to Figure 6.**
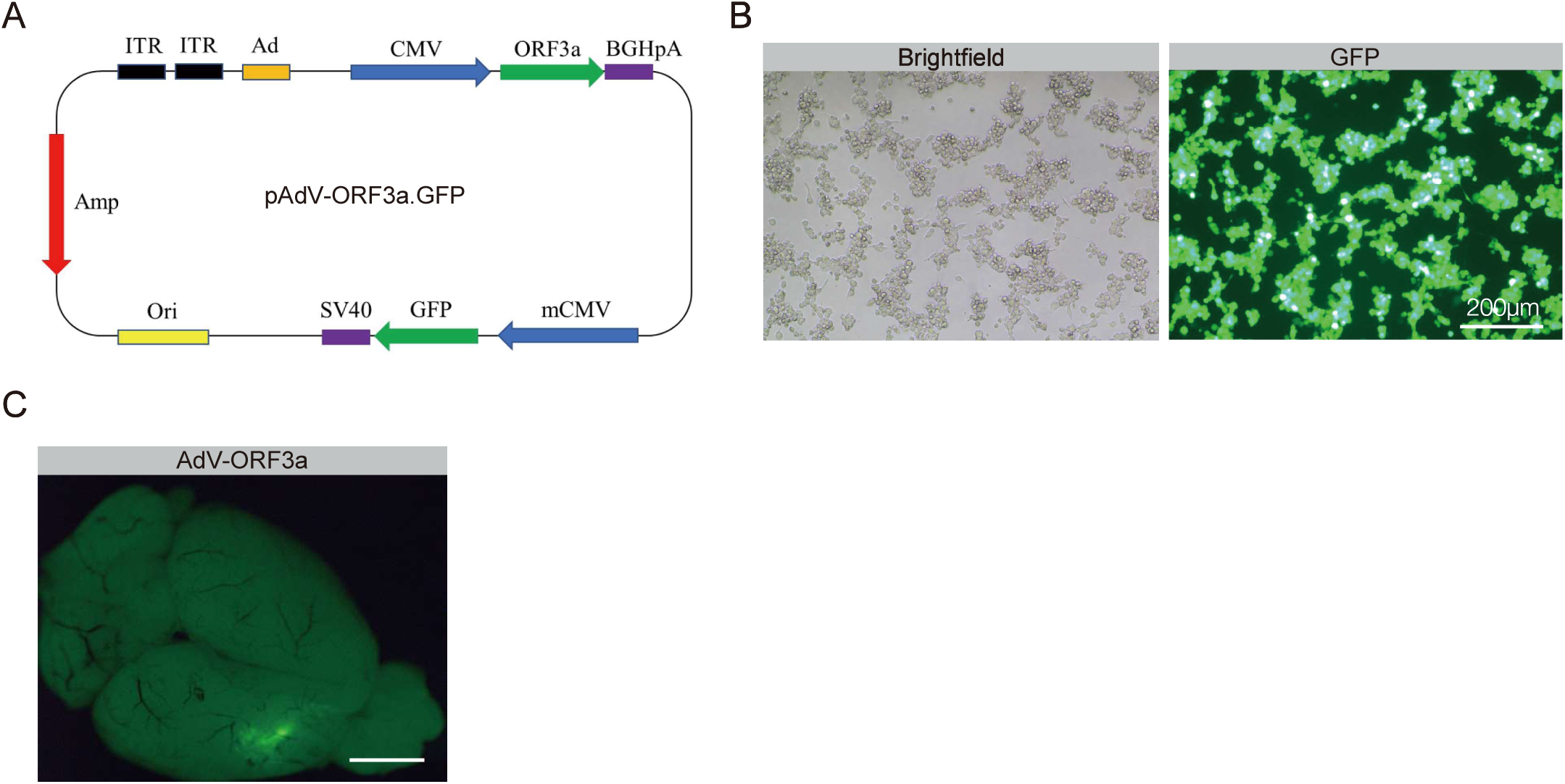
**A.** Vector map of AdV-ORF3a.GFP. **B.** Micrograph of HEK293 cells transduced with AdV-ORF3a.GFP. Transduced cells expressed ORF3a.GFP. **C.** Micrograph of a whole mount brain after retro-orbital injection of AdV-ORF3a.GFP. Transduced neural tissue expressed ORF3a.GFP.

**Figure S6 Related to Figure 7.**
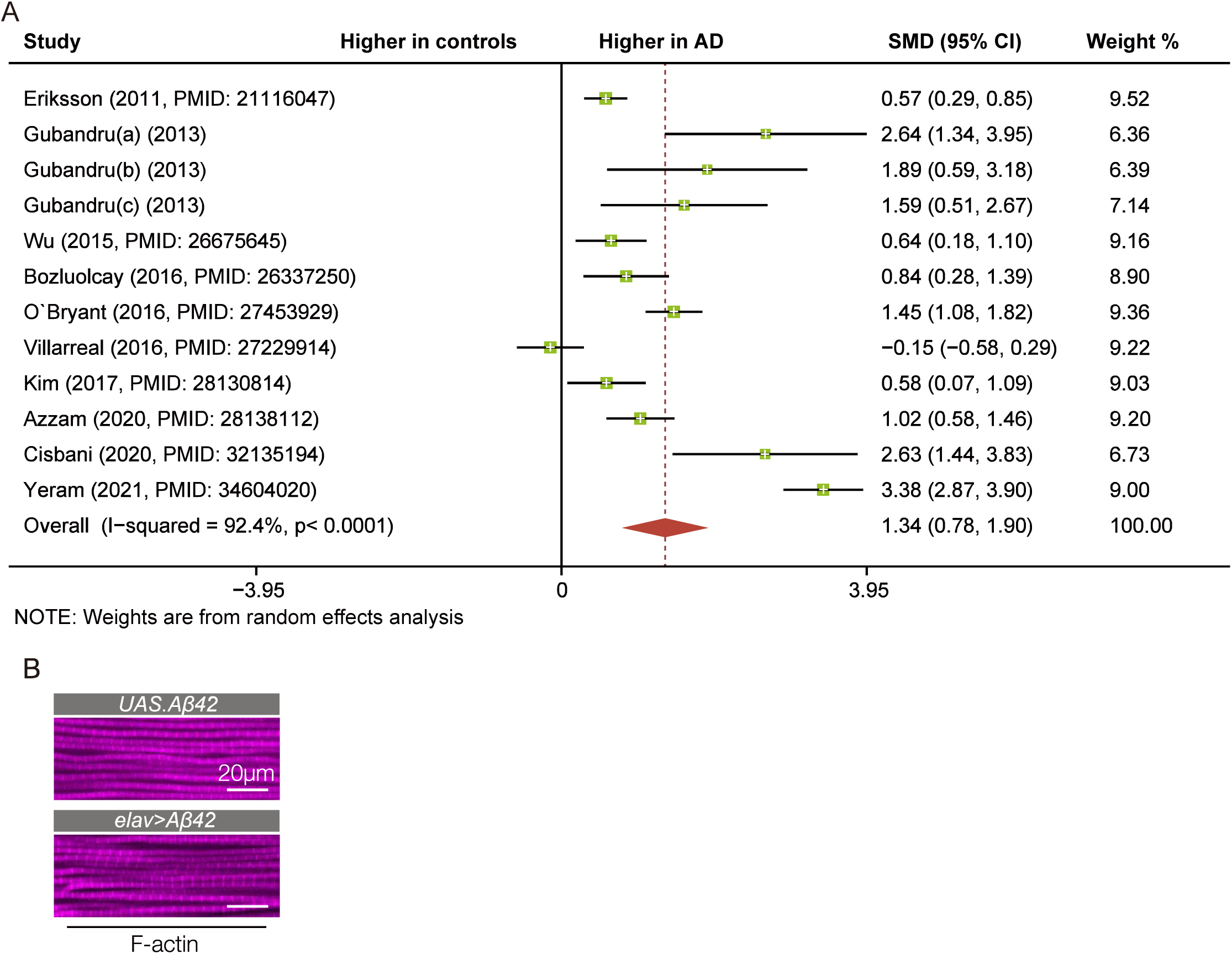
**A.** Forest plot depicting IL-6 levels in AD patients. Squares represent the odds ratio and horizontal lines show the 95% confidence intervals. The solid vertical line corresponds to no effect. The red diamond shows the summary measure, indicating the serum levels of IL-6 were increased in AD patients (n=585) compared to health controls (n=439). **B.** Confocal micrographs of indirect flight muscle stained with phalloidin to visualize F-actin (violet). Muscle morphology was comparable between control (*elav>Gal4*) and *elav>Aβ42* flies.

## Notes

### Competing Interest Statement

The authors have declared no competing interest.

### Summary of Updates

In the revised manuscript, we identify an inter-organ communication pathway that is activated by neuroinflammation. The signaling ligand Upd3/IL-6 is expressed in the brain in response to neuroinflammation, and activates the JAK/Stat signaling pathway in skeletal muscle. Once active, the JAK/Stat pathway inhibits mitochondrial function and reduces skeletal muscle performance. Neuroinflammation therefore activates a systemic signaling response that alters muscle performance.

